# Macrophages break interneuromast cell quiescence by intervening the inhibition of Schwann cells in zebrafish lateral line

**DOI:** 10.1101/2021.12.29.474498

**Authors:** Meng-Ju Lin, Chia-Ming Lee, Wei-Lin Hsu, Bi-Chang Chen, Shyh-Jye Lee

**Author notes:** Department of Plastic Surgery, New York University, 540 First Avenue, Skirball Institute, New York, NY 10016, USA.

## Abstract

In the zebrafish lateral line system, interneuromast cells (INCs) between neuromasts are normally kept quiescent by underlying Schwann cells (SWCs). Upon severe injuries that cause the complete loss of an entire neuromast, INCs can occasionally differentiate into neuromasts but how they escape from the inhibition by SWCs is still unclear. Using a genetic/chemical method to specifically ablate a neuromast, we found a small portion of larvae can regenerate a new neuromast, but the regeneration was hindered by inhibiting macrophages. By *in toto* imaging, we further discovered heterogeneities in macrophage behavior and distribution along lateral line. We witnessed the crawling of macrophages in between injured lateral line and SWCs during regeneration and also in between the second primordium and the first mature lateral line during development. It implies that macrophages may physically separate and alleviate the inhibition from pLLn and SWCs to break the quiescence of INCs during regeneration and development in the zebrafish lateral line.

## INTRODUCTION

Tissue regeneration is a critical step to maintain homeostasis and restore organ function after injury in multicellular organisms. Planarians form neoblasts to replenish the whole organism from finite body parts [1, 2]. In contrast, injury results in massive scarring that hampers regeneration in mammals, so mammals only retain a limited regeneration capacity to replace their damaged tissues. Scars often cause irreversible cell loss and damages to tissues and organs. For example, the destruction of mechanosensory hair cells within inner ear by antibiotics, noise or aging may cause hearing deficit. However, in lower vertebrates like birds, amphibians and fish, hair cells could be robustly and continuously recovered after damage [3–7]. Intriguingly, multipotent progenitor cells residing in mammalian cochlea could be cultured *in vitro* under some conditions that potentially could be used to complement hair cell loss [8–11]. It suggests that genetic machinery for tissue renewal is still retained in most animal species but not the regulatory mechanisms required to awaken the quiescence of stem cells [12, 13]. However, outstanding questions still remain elusive. For examples. how the regeneration is triggered? How the responsiveness of progenitor cells is regulated at the cellular and molecular levels and how the degree of triggers may result and/or regulate differential responsiveness of potential progenitor cells? To investigate these questions, we study zebrafish (*Danio rerio*), a well-established vertebrate model with remarkable regeneration capacity in most tissues and organs, including lateral line system.

The lateral line system is a mechanosensory system that detects water movements around fish body, contributing to navigation, schooling and predator avoidance [14, 15]. The zebrafish lateral line system includes anterior and posterior lateral lines flanking both lateral sides of fish body. The sensory organs of the lateral line system are neuromasts. Taking the posterior lateral line (pLL) as an example, during development neuromasts are periodically deposited during the posterior migration of a pLL primordium (pLLp), which is a migrating cluster of about a hundred cells [16]. A neuromast is composed of centrally-positioned hair cells, which are functionally and structurally similar to the hair cells of mammalian inner ear. Hair cells are protected by surrounded supporting cells or mantle cells (MCs), which are progenitor cells during hair regeneration (Fig. 1A) [17, 18]. The lateral line hair cells are exposed on the body surface and constantly face mechanical and/or chemical environmental assaults. Due to the superficial location, hair cells can be easily ablated in zebrafish by exposing zebrafish larvae to heavy ions like copper [7, 19, 20], mercury [21] or antibiotics such as neomycin [22]. A previous study reported that supporting cells underneath hair cells are notable progenitor cells in contrast to the center or anterior differentiating or dormant support cells [6]. In addition, MCs encircling the neuromast are quiescent cells, however, they can re-enter the S phase to form a new neuromast facing a high copper ion concentration [6] and a severe injury like tail-fin amputation [23, 24]. Lastly, interneuromast cells (INCs) sitting in between neuromasts are also kept quiescent by the underlying Schwann cells (SWCs) and pLL nerve (pLLn) connecting to hair cells. Perturbation of epidermal growth factor (EGF) pathway between SWCs and pLLn results in early activation of INCs and precocious intercalary neuromast formation [25–28]. The intercalary neuromast formation also occurs during the migration of the 2^nd^ pLLp by intervening the contact between INCs and SWCs [18, 29, 30]. It has also been shown that a new neuromast can be regenerated by electro-ablating a whole neuromast [31]. These studies nicely demonstrate the regeneration capacity of INCs. By RNA-seq analysis and functional assay, MCs and INCs have been shown to contribute to hair cell regeneration by interrogating multiple signaling pathways such as Notch, Wnts, Fgfs and retinoic acid (RA) sequentially and spatially [32, 33]. However, it is still puzzling to understand the cellular hierarchy of lateral line regeneration while facing diversified external cues.

**Figure 1.**
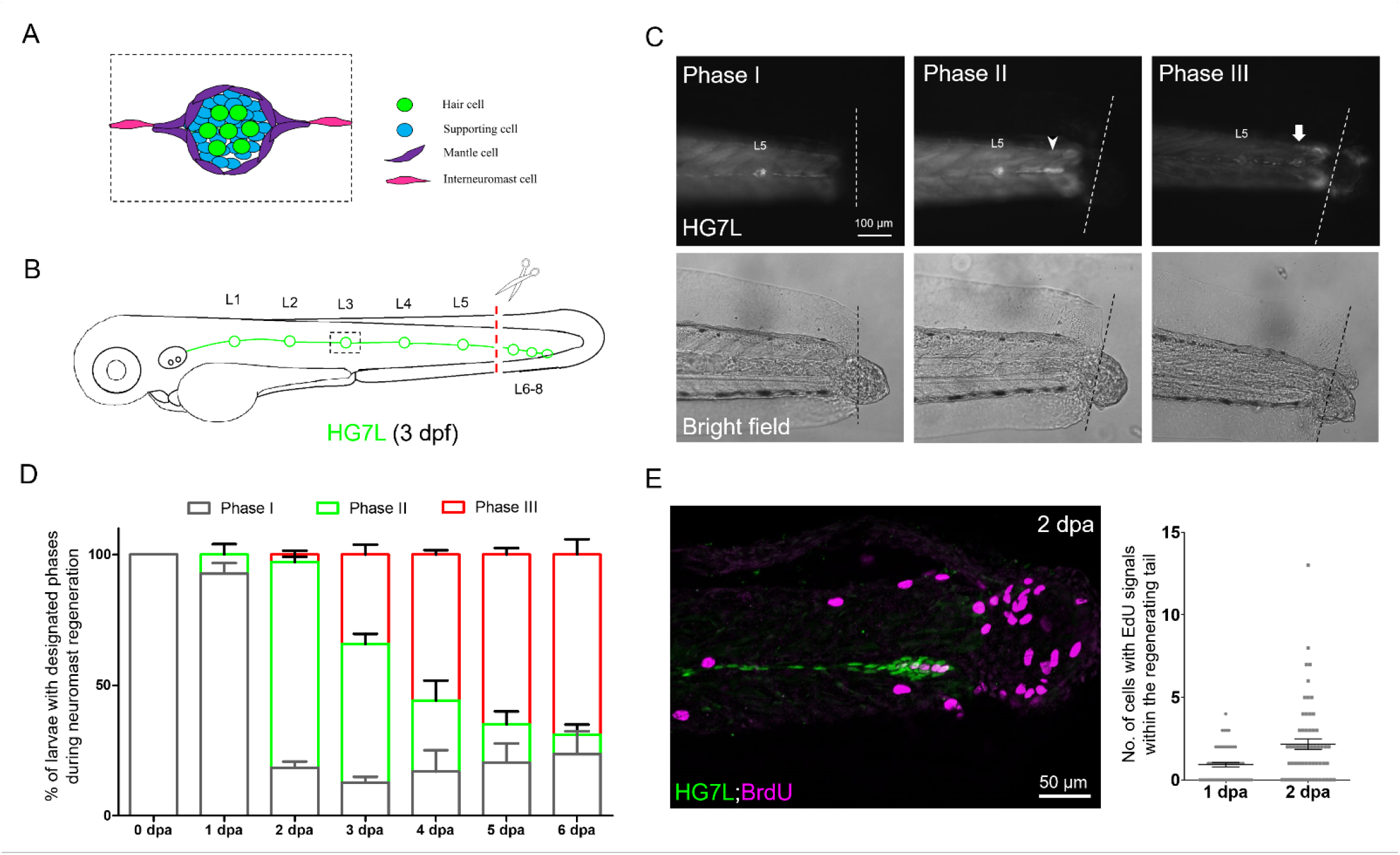
Active cell proliferation and clustering occur during neuromast regeneration upon fin amputation. (A) A cartoon shows different cell types in a neuromast of lateral line. (B) A cartoon shows only one side of the fluorescent posterior lateral (in green) with L1-8 neuromasts of an *Et(HG7L)* larva at 3 days post fertilization. The tail fin is clipped at the dashed line to remove neuromast L6-8. (C) A new neuromast was regenerated in three distinct phases as examined at the GFP channel under epifluorescent microscopy. Phase I: No notable increase in fluorescent cells were observed in the lateral line between the L5 neuromast (as labeled) and the cut site (dotted line). Phase II: Fluorescent cells were increased and aggregated to form a cluster (arrowhead). Phase III: A new neuromast was formed (arrow). The corresponding bright field images for each phase are shown below. (D) The percentages of larvae at each phase were calculated at designated day post amputation (dpa, N = 3, n = 70). (E) HG7L larvae were fin-amputated, fixed at 1 and 2 dpa and subjected BrdU staining (in magenta) to probe actively proliferating cells or GFP immunohistochemistry (in green) to stain lateral line. Active cell proliferation was observed near the cutting edge at 2 dpa and quantified as shown in a scatter plot on the right.

Tissue damage elicited macrophage infiltration has recently been demonstrated to be one of factors promoting regeneration [34, 35]. Macrophages acquire polarity transition from the M1-like macrophages, which are classically regarded as activated pro-inflammatory cells, to the M2-like phenotype known as alternative activated macrophages involved in inflammation resolution [36, 37]. To suppress the chronic inflammatory response, macrophages clean up infected neutrophils by phagocytosis [38] and secret anti-inflammatory cytokines such as IL-10 and TGFβ [39, 40]. Recently, macrophages were found to enhance regeneration by secreting cytokines. Macrophage-secreted TNFα was demonstrated to be critical for blastemal formation during fin regeneration in zebrafish [41]. Angiogenesis induced by the secretion of VEGF from macrophages could also assist in wound repair and regeneration of blood vessels and peripheral nerves [42–44]. Macrophages could also go through either canonical or non-canonical Wnt pathway to influence regeneration [45, 46]. In addition to the effects of cytokine, macrophages could exert mechanical forces to facilitate wound repair and regeneration [42, 47]. As expected, macrophages play a pivotal role in hair cell regeneration in contrast to neutrophils [48]. However, how macrophages work to facilitate lateral line regeneration upon tissue damage remains unclear.

In this work, we first provide further evidences to support that INCs are the major progenitor cell source to replenish an entire neuromast post tail fin amputation. To our surprise, a few larvae were still found to regenerate a neuromast post specific chemical-genetic ablation without harming SWCs and pLLn. By *in toto* imaging, we found macrophages are significantly patrolling at damage sites. In addition, we also observed that macrophages squeeze in between INCs and SWCs/pLLn. It suggests that macrophages break INC quiescence by intervening the inhibition of SWCs in zebrafish lateral line system.

Abbreviations: interneumast cells (INCs), supporting cells (SCs), hair cells (HCs), mantle cells (MCs), Schwann cells (SWCs), days post fertilization (dpf), days post amputation (dpa)

## MATERIALS AND METHODS

### Ethics Statement

All animal handling procedures were approved by the use of laboratory animal committee at National Taiwan University, Taipei, Taiwan (IACUC Approval ID: 107 Animal Use document No. 00180) and were carried out in accordance with approved guidelines.

### Zebrafish strains

Wild type AB zebrafish (*Danio rerio*), *Tg(-4.7alpl:mCherry)* [32], *Tg(-8.0cldnb:lyn-GFP)* [49], *Tg(FoxD3:GFP)* [50], *Tg(mpeg1:mCherry)* [51], *Tg(mpx:GFP)* [52] and *Tg(6XTcf/LefBS-miniP:d2GFP)^isio1^* [53] fish were maintained at 28.5°C on a 14-h light/10-h dark cycle. To generate the *Et(HG7L)* line, a transgenic cassette was generated by combining p5E-*hsp70*, pME-EGFP and p3E-polyA into pDestTol2CG2 via Gateway recombination cloning [54]. This hsp70-EGFP vector (HG) was proved to be suitable for enhancer trap screen (Fig. S1A) [55]. Therefore, we conducted an enhancer trap line screen and identified a trap line, *Et(HG7L)*, which was named after its specific expression in the pLL system (Fig. S1G-M), and was utilized in this study. To generate the *Tg(-8.0cldnb:NTR-hKikGR; myl7:egfp)* line, we generated a cassette for transgenesis by combining p5E-*8.0cldnb* (a kind gift from Dr. Tatjana Piotrowski) [6], pME-NTR-hKikGR (a kind gift from Dr. Chung-Der Hsiao) [56] and p3E-polyA into pDestTol2CG2 through Gateway cloning. For transgenesis, both Tol2 transposon and Tol2 transposase mRNA were injected at 25 pg into 1-2 cell stage embryos and raised to adults (F0). We backcrossed F0 founder fish with wildtype fish, screened F1 embryos with strong EGFP signals in heart and selected one founder with a strong expression in the pLL system as shown in Fig. S3. We also generated double transgenic lines, including *Tg(-8.0cldnb:NTR-hKikGR); Et(HG7L)*, *Tg(mpeg1:mCherry; FoxD3:GFP)*, *Tg(mpeg1:mCherry; -8.0cldnb:NTR-hKikGR)*, *Tg(-4.7alpl:mCherry; 6XTcf/LefBS-miniP:d2GFP) and Tg(mpeg1:mCherry; mpx:GFP)*, to be used for producing quadruple transgenic larvae as indicated. Embryos collected from natural mating were cultured and staged according to Kimmel et al. [57].

### Whole mount *in situ* hybridization

DNA fragments of *sorcs3* and *ccdc147* was cloned from zebrafish cDNAs by RT-PCR and subcloned into pGEMT-easy vectors for antisense probe synthesis. Primer pairs used are as following: *sorcs3* (forward: GTCGCCAATGCAAGTGAATTACGC; reverse: TTTCCAGACCAGTACACGACTGCGT) and *ccdc147* (forward: GACGACAGTACGTTGGAAACCATGG; reverse: CGGTGGCTTTAGTAAGGTTTTCCCG). Whole-mount *in situ* hybridization (WISH) was performed as described using digoxigenin (DIG)-labeled antisense RNA probe [58]. Stained embryos were mounted in glycerol, observed under a Leica S8AP0 stereomicroscope (Leica Microsystems, Wetzlar, Germany) and photographed using a Canon 7D DSLR camera (Canon, Tokyo, Japan).

### Immunohistochemistry and confocal microscopy

Whole-mount immunohistochemistry (IHC) staining was performed as previously described [59] by using either mouse anti-GFP antibody (GT859, GeneTex, 1:500), rabbit anti-histone H3S10ph (phosphor Ser10) antibody (GTX128116, GeneTex, 1:1000), mouse anti-ZO-1/TJP1 antibody (33-9100, Thermo Scientific, 1:100), rabbit anti-GFP antibody (GTX113617, GeneTex, 1:500), mouse anti-tubulin (acetyl Lys40) antibody (32-2700, Thermo Scientific, 1:1000) and rat anti-mCherry antibody (M11217, Molecular probes, Thermo Scientific, 1:300). Secondary antibodies used are as following: goat anti-mouse or anti-rabbit IgG conjugated with Alexa Flour 488 or Alexa Fluor 568 (Molecular probes, 1:500). Confocal images were collected utilizing LSM 780 or LSM 880 confocal laser-scanning microscope with 20X lens or 43X water lens (Carl Zeiss, Oberkochen, Germany). In general, 8 to 20 layers with 1 μm thickness were scanned and stacked for each image unless otherwise stated, further processed and presented as maximum intensity projection by the Fiji software [60].

### Cell proliferation analysis and TUNEL staining

To detect proliferating cells, embryos were first treated with 10 mM 5-bromo-2’-deoxyuridine (BrdU, Sigma-Aldrich) pulses at designated stage and then fixed with fresh 4 % paraformaldehyde (PFA) in phosphate-buffered saline (PBS). Whole-mount IHC was performed with mouse BrdU antibody (B2531, Sigma-Aldrich, 1:250). Apoptotic cells were also examined by TUNEL assay. Embryos were fixed with 4% PFA/PBS overnight at 4°C and dehydrated with methanol at −20°C. After gradual rehydration, the embryos were permeabilized with 10 µg/ml proteinase K for 2 min at room temperature then post-fixed by 4% PFA/PBS. After several washes of PBS-T (0.1% tween-20 in PBS), embryos were again fixed by pre-chilled solution containing ethanol and acetic acid (in 2:1 ratio) at −20°C for 10 min. After the pH value was brought back with washes of PBS-T, samples were incubated with 27 µl labeling solution plus 3 µl enzyme solution (In Situ Cell Death Detection Kit, AP, Roche) at 37°C overnight. They were washed three times with PBS-T for 5 min each and were further performed with double whole-mount IHC staining with mouse anti-tubulin (acetyl Lys40) antibody and rabbit anti-GFP antibody.

### Time-lapse movies with light sheet fluorescence microscope (LSFM)

The time lapse movies revealing the morphogenesis of neuromast regeneration were taken by the Zeiss Lightsheet Z. 1 (Carl Zeiss, Oberkochen, Germany). The other recordings showing macrophages dynamics were taken by a simple light sheet platform built by Dr. Bi-Chang Chen. This platform fascinatingly features (1) Bessel beam scanning, (2) dual illumination arms, (3) multiple software compatibility (μManager), (4) flexible objective combination, (4) large chamber as 12 mm×12 mm×25 mm, and (5) long-range xyz motorized stage (MS-2000, Applied Scientific Instrumentation, USA).

### Tracking analysis with plugin “TrackMate”

To track the infiltration of macrophage during regeneration, we utilized a plugin “TrackMate” in the Fiji software [61]. Since the z-sections focused were thin, we simplified the context by projecting z section with maximum intensity. To detect spots of interest, we selected LoG (Laplacian of Gaussian) detector with an estimated bulb diameter of 30 pixels. We used simple LAP tracker with maximum linking distance (50 pixels), gap-closing maximum distance (15 pixels) and gap-closing maximum frame gap (3 frame gaps) to generate tracking paths. All spots and links were manually corrected. Filters (Y position and Quality) were used to exclude spots which did not interact with the pLL system or at low quality. To display, spots were labeled with radius ratio of 0.3∼0.4 and tracks were presented with a color map set by Track index.

### Nitroreductase (NTR)/Metronidazole-mediated neuromast ablation

Embryos from the cross of *Et(HG7L)* and *Tg(-8.0cldnb:NTR-hKikGR)* were collected and screened for double positive ones. Freshly-made 10X stock of metronidazole (Mtz, M3761, SIGMA-ALDRICH) solution (20 mM Mtz, 1 % DMSO) were prepared. Larvae at 3 days post fertilization (dpf) were first treated with 1X working concentration of Mtz solution (2 mM Mtz, 0.1 % DMSO diluted in 0.3 % PTU containing 0.3X Danieau’s buffer) for 3, 6, 9, 12 hours in the dark. Then, Mtz solution was replaced with several washes of fresh 0.3X Danieau’s buffer for recovery. The treatment of Mtz solution was 12 hours to obtain the optimal ablation result.

### Pharmacological inhibitors

All the chemical inhibitors were diluted in distilled water except for AG1478 (T4182, SIGMA-ALDRICH, in dimethyl sulfoxide (DMSO)) and LMT-28 (SML1628, SIGMA-ALDRICH, in methanol). Therefore, we used either 0.1 % DMSO or 0.1 % methanol as negative controls of AG1478 or LMT-28 treatments. AG1478 was added at 3 μM to the medium right after tail amputation. To disrupt different signaling pathways during Mtz ablation, we treated 3 dpf larvae with designated drugs together with Mtz. After neuromast ablation, the chemical inhibitors were either washed out with Mtz or added back to be kept in the medium. The COX inhibitor Diclofenac sodium salt (D6899, SIGMA-ALDRICH), a specific blocker of neutrophil recruitment, was used at 3 μM. The TNFα inhibitor pentoxifylline (PTX, P1784, SIGMA-ALDRICH) was used at 35 μM. The IL-6 inhibitors LMT-28 and the Wnt/β-catenin inhibitor IWR-1 (I0161, SIGMA-ALDRICH) were both used at 10 μM.

### Liposome Injection for macrophage ablation

To kill macrophages, 3-dpf larvae were anesthetized with 0.016 % Tricaine/Ethyl 3-aminobenzoate methanesulfonate salt (A5040, SIGMA-ALDRICH) and microinjected with 5-8 nL of liposome encapsulated clodronate (SKU# CLD-8909, Clodrosome) into the circulation system via the Duct of Cuvier. The injection needle was prepared using borosilicate glass microcapillary needles (Sutter Instrument CO., B100-75-10, 1 mm O.D. X 0.75 mm I.D., Novato, CA, USA) with a Sutter puller (P-97, Sutter Instrument) with the following settings for shorter tip: air pressure 500, heat 510, pull 100, velocity 200, time 60 [62]. Control liposomes supplied in the kit were used as controls.

### Statistical analysis with R software

All experimental values except Fig. 7 are presented as mean ± standard error and were analyzed by one-way ANOVA. The number in bottom or above the bar indicates the total sample number in one experimental condition. Groups denoted with different lettering refer to statistical significance (p < 0.05). In Fig. 7, raw data obtained from TrackMate plugin, including spots, links and tracks information were further processed, analyzed and presented as scatter plot or histogram by “ggplot2” package in the R software.

## RESULTS

### Regeneration of neuromast post fin amputation

To investigate how the pLL system regenerates an entire neuromast, we removed the distal neuromast cluster (L6-8) by amputating the caudal fin of a larva from an enhancer-trap line, *Et(HG7L)*, which expresses strong enhanced green fluorescent protein (eGFP) in both MCs and INCs in the lateral line (Supplementary Fig. S1), at 3 days post fertilization (dpf) as indicated in Fig. 1B. We observed the change of fluorescent lateral line at designated time points in bright and dark fields under an epi-fluorescent microscope. A significant portion of larvae grew a new neuromast near the wound site within days. We divided the entire process into three phases. Phase I: no cell clustering; Phase II: cell clustering; Phase III: formation of a new neuromast (Fig. 1C). At one day post fin amputation (dpa), the majority of larvae (92.7%, n = 70) stayed at the Phase I. However, they quickly shifted to the Phase II in 78.7% of larvae at 2 dpa. The formation of cluster might be due to active cell proliferation, and indeed cell proliferation at the lateral line was significantly increased as examined by BrdU labeling at 2 dpa compared to that of 1 dpa (Fig. 1E). Even though no cell clustering was notable at Phase I, cell proliferation did occur near the cut site of the lateral line. It suggests that cells gradually pile up to form a cluster as seen in Phase II.

A rosette appears in the center of a developing neuromast [17]. We also observed the formation of rosette as evident with an intensified eGFP signal (Fig. S2C, arrowheads) in the center of a regenerating cluster of a fin-amputated larva from the cross of *Tg(-8.0cldnb:lyn-GFP)* and *Tg(-4.7alpl:mCherry)* lines (Fig. S2B) [49, 63]. Tight junctions are formed during apical cell assembly in the rosette [17]. We, therefore, confirmed the existence of rosette by immunostaining against ZO-1, a tight junction associated protein [63], in 2-dpa *Et(HG7L)* larvae within the regenerating cluster (Fig. S2D). Interestingly, the cluster requires a larger area (1715.4 μm^2^, n = 10) and a smaller length/width ratio (6.79, n = 10) to accommodate one rosette, with its center mostly situated in the middle of cluster (n = 16) (Fig. S2E). At 4 dpa, more than half of larvae (55.8 %) had regenerated a new neuromast during observation.

Next, we examined the regeneration process more thoroughly under light-sheet fluorescence microscopy (LSFM), which is less photo-toxic and allows long-term recording [64]. Multiple active cell protrusions (arrowheads) were observed in the leading end of pLL system of larvae at both Phase I & II stages (Supplementary Video 1 & 2). Unlike traditional collective migrations such as the pLLp migration during development, new cells climbed onto the original pLL system. They migrated toward injury sites at Phase I and visible protrusions were found at the lagging end of cluster at Phase II. Afterwards, the homogenous cell cluster transformed into polarized cells with ring-like features of MCs within a new neuromast (Supplementary Video 3). Altogether, a new neuromast is regenerated by active cell proliferation, clustering and stochastic cell migrations that is dissimilar to typical cell movements during pLLp development.

### Interneuromast cell is the major cell type contributing to neuromast regeneration post fin amputation

It is well known the loss of neuromast hair cells is replenished by underlying supporting cells (SCs) [65] driven by differential Wnt or Notch signaling [6]. In contrast, the roles of MCs and INCs are less appreciated. MCs seem to stay in quiescence unless facing severe damage [6]. Given that different injury levels may arouse differential regeneration responses of the pLL system, we examined which type(s) of neuromast progenitor cell is/are the major progenitor(s) for regenerating neuromasts post fin amputation. Since the *Et(HG7L)* line also expresses weak eGFP in a portion of SCs, we first used their larvae crossing with *Tg(-4.7alpl:mCherry)* lines with a dim mCherry (red) fluorescence found in only MCs and INCs (Fig. S1I-M). It appeared that the clusters are mainly showing overlapping signals in the regenerating cluster at 2 dpa (Fig. 2A). Moreover, we crossed the *Et(HG7L)* line with the *Tg(-8.0cldnb:lyn-GFP)* line expressing strong membrane-bound eGFP in all neuromast cell types and found that the clustering cells showed both membrane and cytosol eGFP in all cells (Fig. 2B).These both suggest that the clustering cells are mainly from MCs and INCs but not SCs.

**Figure 2.**
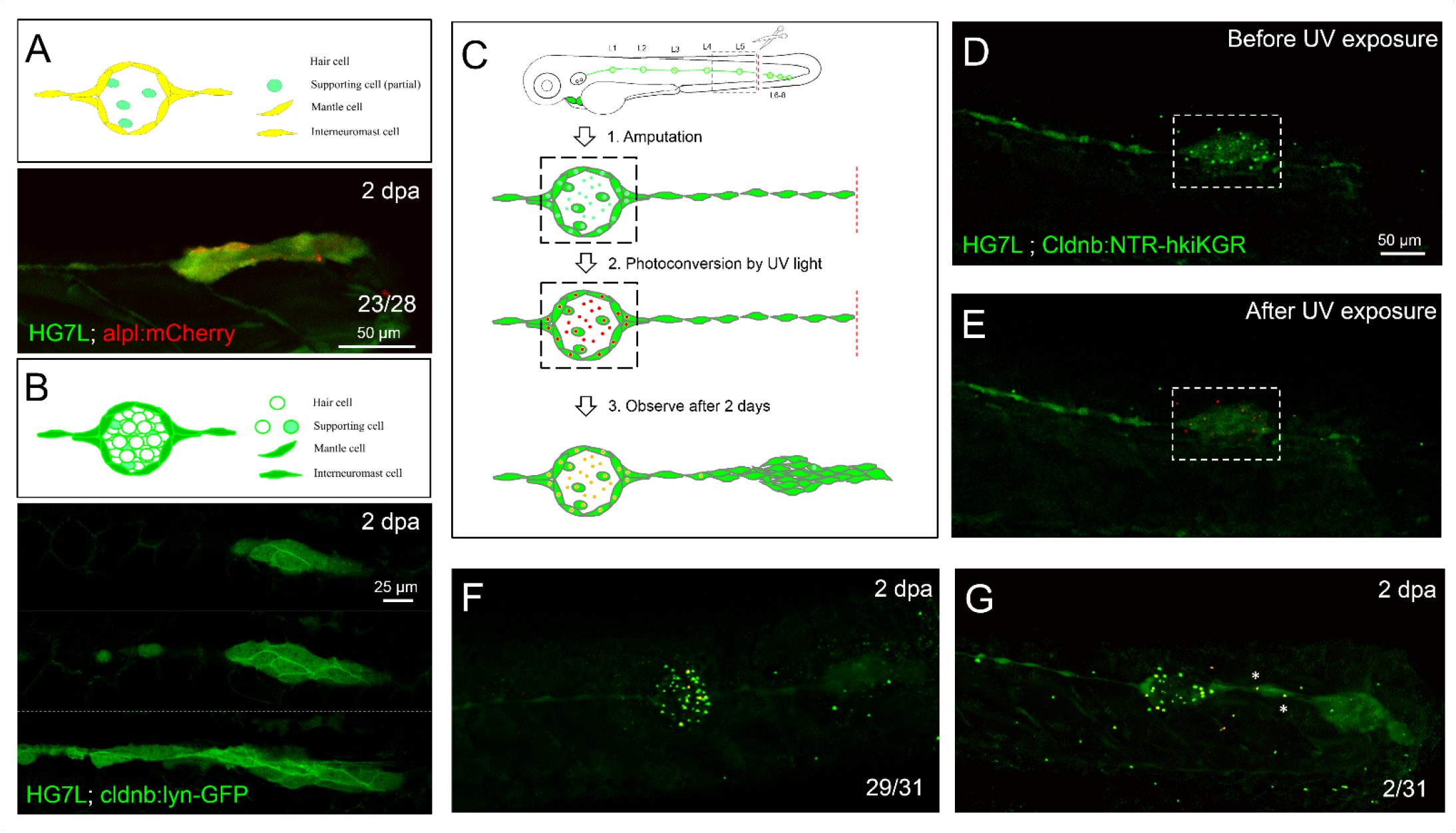
Interneuromast cells are the origin of clustering cells post fin amputation. We performed fin amputation on double transgenic larvae from the cross of the *Et(HG7L)* with the *Tg(-4.7alpl:mCherry)* (alpl:mCherry) or *Tg(-8.0cldnb:lyn-GFP)* (cldnb:lynGFP) to track the origin of progenitor cells for the regenerating cluster post fin amputation at two days after amputation (2 dpa). (A) The regenerating cluster of fin amputated-larvae of *Et(HG7L)XTg(-4.7alpl:mCherry)* lines was examined at GFP or mCherry channel and photographed under confocal microscopy. A representative superimposed image from both channels is shown. The regenerating cluster image contains mostly yellow signal. As illustrated in a cartoon above, the clustering cells could be originated from both interneumast cells (INCs) and mantle cells (MCs). (B) The regenerating cluster of fin amputated-larvae of *Et(HG7L)* X *Tg(-8.0cldnb:lyn-GFP)* was examined at GFP channel to scan the z-axis. Images at three different z positions clearly show the existence of both membrane and cytosol green fluorescence. As illustrated in a cartoon above, the signals could be originated from INCs, MCs and supporting cells. (C) A series of cartoons show the scheme of utilizing a *Tg(-8.0cldnb:NTR-hKikGR; myl7:EGFP)* transgenic line, which expresses nitroreductase (NTR) and hKikGR fusion protein, to further discern whether the clustering cells are from INCs or cells inside a neuromast. To label the lateral line, we used larvae from the cross of *Et(HG7L)* (green) and *Tg(-8.0cldnb:NTR-hKikGR; myl7:EGFP)* (green dots), examined and photographed at GFP channel before (D) and after UV exposure (E). After UV exposure, the green *hKikGR* dots were converted to red, in all cell types including MCs within distal neuromast right after fin amputation. (F) At 2 dpa, labeled cells generated new green hKikGR protein. The mingling of green and red signals become yellow as shown in (F). The labeled cells within the neuromasts rarely escaped in most treated larvae (29/31). However, we observed that a few larvae (2/31) showed yellow signal in the neighboring of INCs (white asterisks) as shown in (G).

Since none of *in vivo* labeling method or transgenic line is available to distinguish MCs and INCs for their roles in neuromast regeneration, we established a double transgenic line, *Tg(-8.0cldnb:NTR-hKikGR; myl7:egfp)*, with a *claudin-b* promoter driven expression of a fusion protein, of nitroreductase (NTR) [66] and a photo-convertible fluorescent protein (humanized Kikume Green-Red, abbreviated as hKikGR hereafter) [67] in all cell types of the pLL system (see gene constructs in Fig. S3A). A *myl7*-driven eGFP gene was also included for making fluorescent heart as a selection marker. The NTR-driven cell ablation will not be mentioned until later experiments. Here, the hKikGR protein was expressed in punctate [56] in all cell types within neuromasts of the entire lateral line system, including anterior and posterior lateral lines (Fig. S3B-C).

We amputated tail fins of 3-dpf larvae from the cross of *Tg(-8.0cldnb:NTR-hKikGR)* and *Et(HG7L)* lines and then photo-converted hKikGR protein within the distal neuromast into red signals by ultraviolet light (Fig. 2C-E, dashed rectangles). Two days after amputation, we found red-fluorescent cells stay in the original neuromast in most larvae (Fig. 2F). Only two out of 31 larvae had a few red-fluorescent cells outside the original neuromast (Fig. 2G, asterisks). It further suggests that INCs, but not SCs nor MCs are the major progenitor cell type to regenerate new neuromasts in the condition tested.

### Fin amputation weakens lateral line nerve signal and upregulates interneuromast cell Wnt activity

To understand how INCs are aroused from a quiescent state (Fig. 3A), we first investigated whether pLLn or SWC was affected after tail amputation due to the reported inhibition of Wnt signaling by epidermal growth factor receptor (EGFR) pathway between SWC and pLLn [26]. We collected larvae from the cross of *Tg(-4.7alpl:mCherry)* and *Tg(FoxD3:GFP)* [50] to reveal the pLL system and SWCs, respectively. Larvae were fixed at 2 hours post amputation (hpa), 6 hpa, 1 dpa and 2 dpa, and then subjected to immunohistochemistry against acetyl tubulin to reveal the pLLn. Although we expected to see possible retraction of injured nerves [68], unexpectedly, both pLLn and SWCs appeared to stay in parallel to the distal end of lateral line (Fig. 3B, dashed lines with arrowheads indicated). However, we found that the fluorescent intensity of pLLn but not SWCs was reduced (at least 60 % compared to proximal end) approaching the distal end (Fig. 3B, left panels) by quantifying averaged intensity at the GFP channel (Fig. 3B, dashed brackets) every 20 μm from the cutting edge. The reduction of signals from 6-hpa (orange curve) and 1-dpa larvae (yellow curve) obviously showed a steeper slope (Fig. 3C) compared to those at 2 hpa or 2 dpa (Fig. 3C, red and green curves, respectively). The reduced pLLn signals implied a potential local diminishment of *neuregulin 1* (*nrg1*) within pLL nerves which may alleviate Wnt inactivation in distal INCs [26].

**Figure 3.**
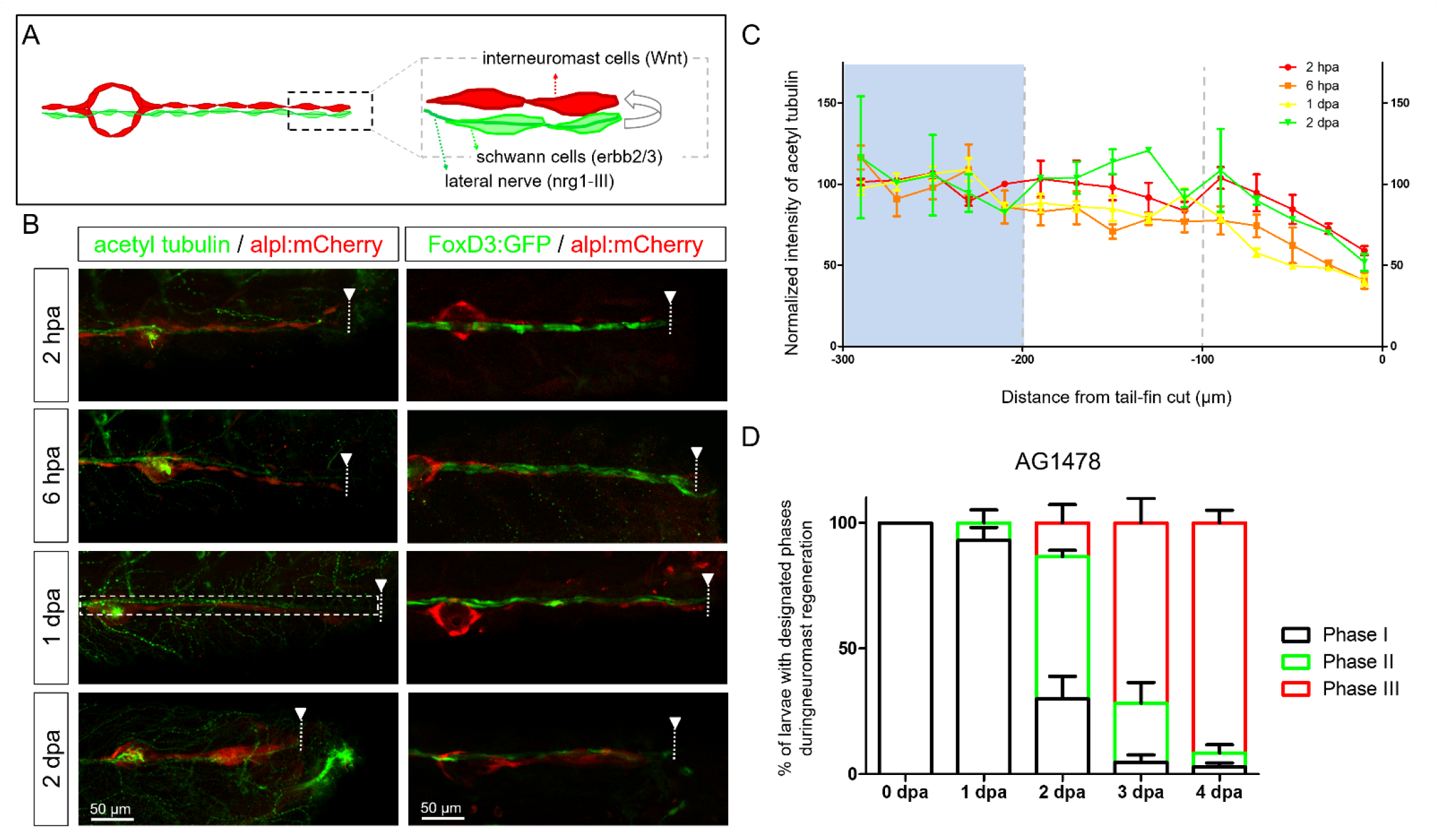
Diminished nerve inhibition near the cut site may allow neuromast regeneration from interneuromast cells. (A) A cartoon shows the close contact between interneuromast cells (INCs, red) with underlying Schwann cells (SWCs, light green) and posterior lateral nerve (pLLn, dark green). The pLLn-derived nrg1-III can activate erbb2/3 receptors on SWCs to suppress Wnt activity in INCs, thus keeps INCs in a quiescent state. (B) The 3-dpf larvae from the cross of *Tg(-4.7alpl:mCherry)* and *Tg(foxd3:EGFP)* were tail fin amputated and immobilized at designated time to show the integrity of SWCs in green (labeled by *Tg(FoxD3:GFP)* in all stages examined (right column). *Tg(-4.7alpl:mCherry)* larvae with red fluorescent lateral line at 3 dpf were also tail fin-amputated, fixed at designated hours or days post amputation (hpa or dpa, respectively) and subjected to immunostaining against acetyl tubulin to reveal green nerves along the red lateral line (left column). It appeared that the fluorescent intensity of green pLLn is diminishing approaching the cut site (dashed line). So, we measured the green fluorescent intensity at decreasing distance from the cut site in cropped images as depicted by a dotted rectangle as shown in the image of 1-dpa larva. The averaged intensity was calculated and normalized with those of the proximal region (200-300 μm from cutting edge, blue background) for designated times shown by different colors. (D)The *Et(HG7L)* larvae at 3 day post fertilization (dpf) were treated with 3 μM of AG1478, fin amputated to examine neuromast regeneration according to Fig. 1C. Data is presented as mean ± s.e.m..

We then blocked EGFR pathway by a EGFR tyrosine kinase inhibitor, AG1478, post fin amputation [69]. The AG1487-treated larvae appeared to have faster recovery of new neuromasts at 2 dpa (Fig. 3D, 13.5 %, n = 57) compared to untreated fish (Fig. 1D, 2.9%, n = 70). Notably, almost all larvae (Fig. 3D, 91.6 %, n = 48) accomplished neuromast regeneration at 4 dpa. To have a more detail observation in real time, we observed the neuromast regeneration process under LSFM. Astonishingly vigorous cell divisions were found (Supplementary Video 4, cell before mitosis labeled by white asterisk, daughter cells marked by magenta asterisks). Besides, considerable number of cell protrusions in various directions were observed (Supplementary Video 4, arrowheads). It suggests that pLLn and SWC play a dominant role in modulating the progenitor status of INCs not only during development [25, 26] but also regeneration.

Since SWCs maintained INCs in a quiescent progenitor by inhibiting Wnt/β-catenin signaling, we next asked whether Wnt reactivation occurs within the pLL system during regeneration. By crossing a Wnt reporter transgenic fish, *Tg(6XTcf/Lef-miniP:d2GFP)* [53] to the *Tg(-4.7alpl:mCherry)* line, we noticed a subpopulation of INCs presenting d2GFP protein (degenerated form) at the distal end from 1 dpa (Fig. 4A-B, n = 17/33). From the close-up view (dashed rectangles with enlarged and split channels shown on right hand side), these subpopulations seemed to correspond with emerging cells which appeared besides existing ones, climbed up and moved toward the injury site (Supplementary Video 1, arrows). Moreover, while cluster appeared at 2 dpa, the regime of Wnt activity expanded as expected (Fig. 4C-D, n = 33/37), with a significant increase in area and range (defined as the distance between proximal and distal end of d2GFP signals) (Fig. 4E-F). These observations suggest that the elevation of Wnt activity within INCs post fin amputation (Fig. 4G).

**Figure 4.**
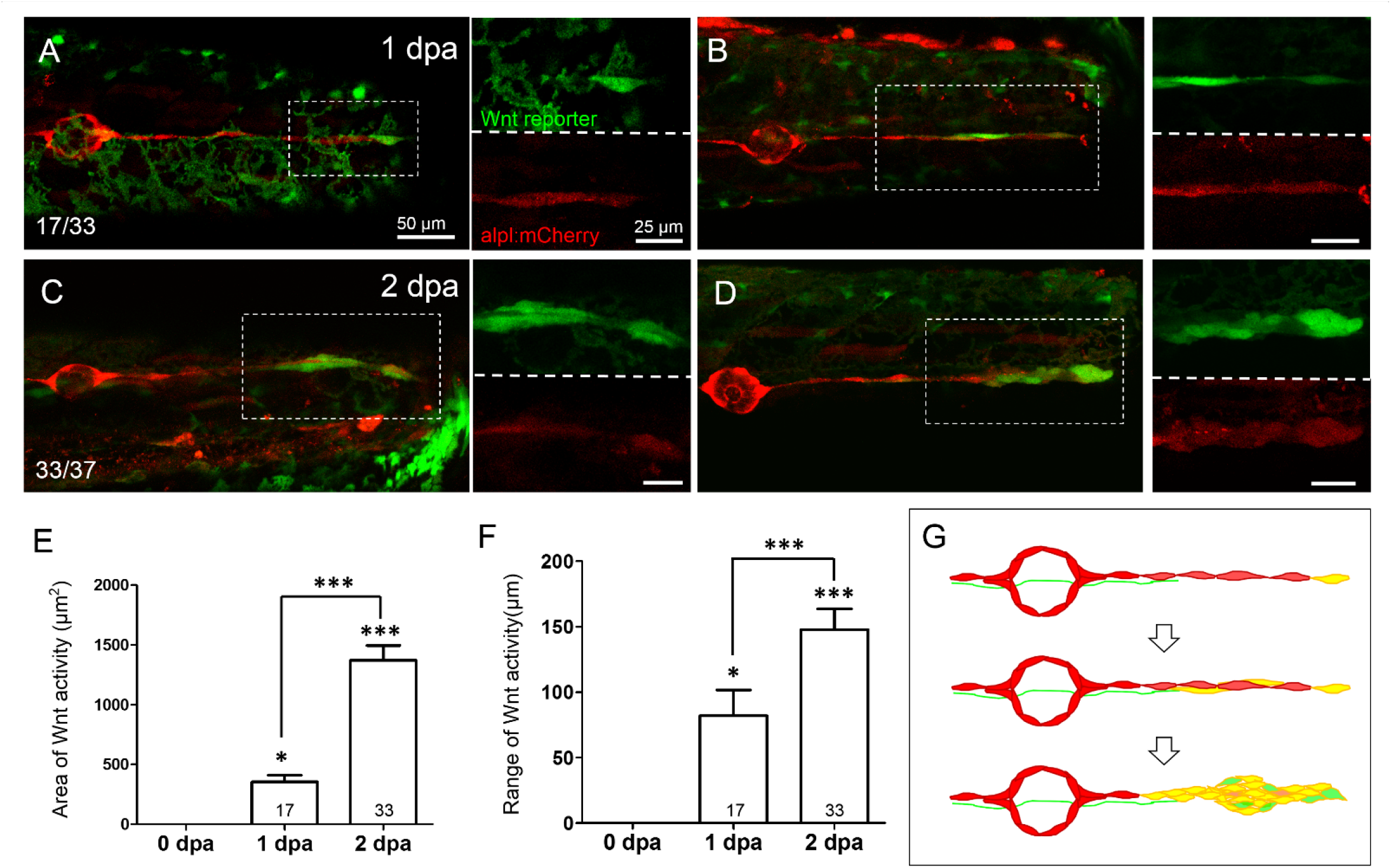
Wnt activity is enhanced in the lateral line near the injured site at 1-2-day post amputation. Representative superimposed stacked images of two different larvae (A,C and B,D) from the cross of *Tg(6XTcf/LefBS-miniP:d2GFP)* (green), and *Tg(-4.7alpl:mCherry)* (red) at 1 day post amputation (dpa, A,B) and 2 dpa (C,D) are shown. Elevated green Wnt activity was observed in and surrounding lateral line as shown in (A). Interneuromast cells (arrows) near the wound site showed yellow signal due to the superimposed color for red lateral line and green Wnt signal. Magnified diagrams for each dashed-rectangle are shown next to the corresponding superimposed photo at green or red channel. Scales are the same for all superimposed images as shown in (A). Scale bars are 25 μm in magnified images. (E-F) The areas and range of Wnt activity, defined by measuring the extension from tail-cut to the most proximal end, were calculated and shown. Data represents mean ± s.e.m. and analyzed by one-way ANOVA comparing to that of 0 dpa. In addition, the difference between 1 and 2 dpa were analyzed and shown. \**P*<0.05, \*\*\**P*<0.0005. (G) A series of cartoons illustrate the elevation of Wnt activity (yellow) during cluster formation.

### Activation of interneuromast cells even in the presence of intact Schwann cells and lateral nerves

A whole neuromast could be destroyed by 100 μM copper sulfate without harming the nearby nervous system [7, 20]. Under this somehow “more” specific ablation, no regeneration of neuromast was observed [31]. It supports the overwhelming inhibition signals coming from pLLn and SWC on the regeneration capacity of INCs. To further test this hypothesis, we used the *Tg(-8.0cldnb:NTR-hKikGR)* line, which specifically expresses NTR in lateral line (Fig. S3). More importantly, we identified a founder fish, which expresses stronger NTR-hKikGR signals in neuromasts compared to that of INCs (Fig. S3B). This line allowed us to chemically kill neuromasts without damaging INCs. To validate this chemical ablation method, we crossed this fish with *Et(HG7L)* to reveal the pLL system and then treated the larvae with metranidazole (Mtz) at 3 dpf. The green fluorescence in neuromast cells was gradually abolished while that of INCs remained unchanged within a half-day incubation of Mtz (Fig. 5B-D). The loss of fluorescence implied the death of cells. By TUNEL assay, we found that cell apoptosis only occurred within neuromasts but not INCs. In addition, no apoptotic signal was found in SWCs and the pLLn in Mtz-treated larvae (Fig. 5F-G). Therefore, this NTR/Mtz ablation system appears to be a convincing approach to specifically kill cells in neuromasts without affecting nearby cells.

**Figure 5.**
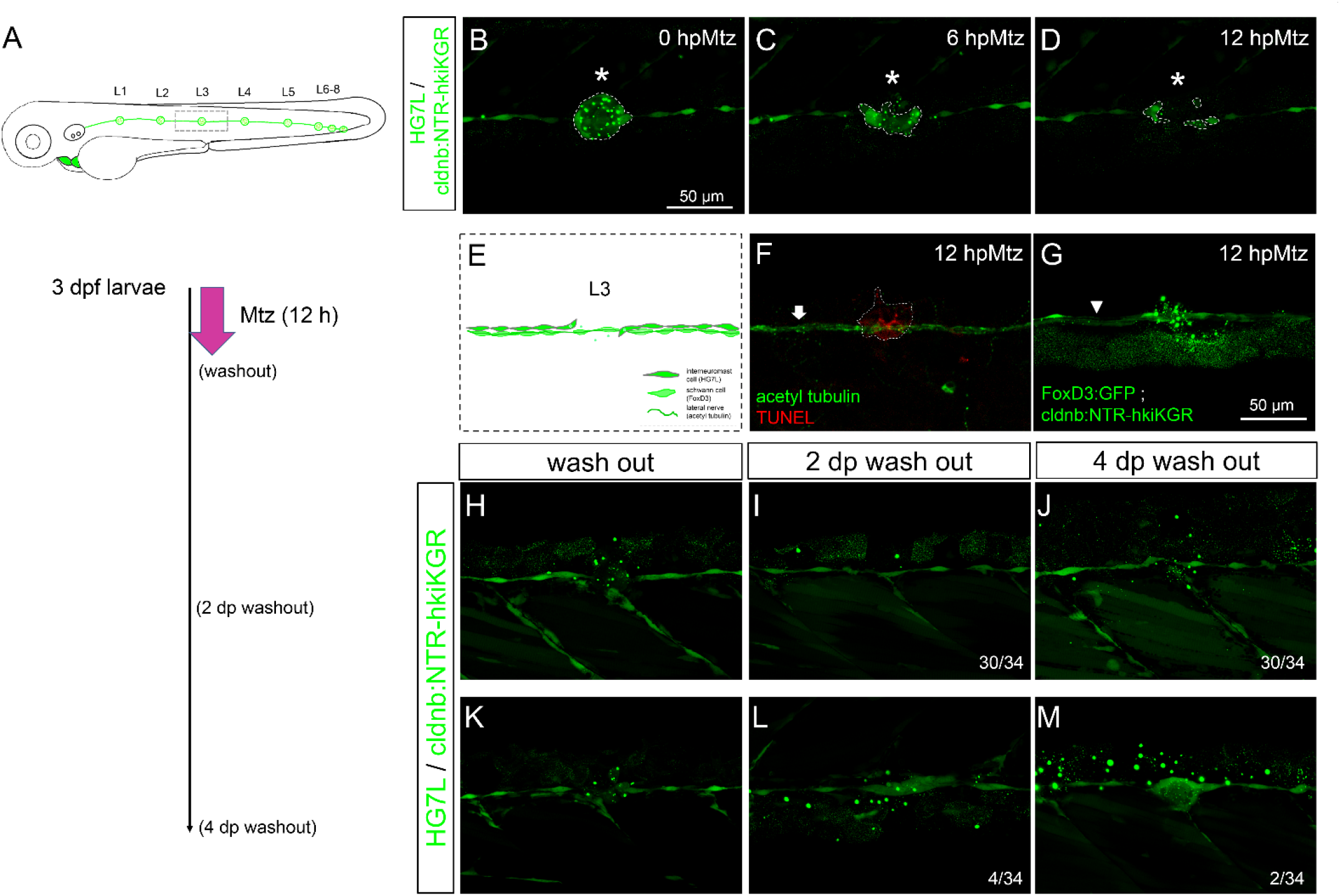
Chemical-genetic ablation of neuromasts blocks neuormast regeneration in most but not all larvae. (A) Larvae from the cross of *Et(HG7L)* (green) and *Tg(-8.0cldnb:NTR-hkiKGR)* (green dots) were treated with 2 mM metronidazole (Mtz) at 3 days post fertilization (dpf) for 12 h, washed out, cultured, examined and photographed at designated h post Mtz treatment (hpMtz) as shown in a series of cartoons. (B-D) Representative stacked images show decreasing size (enclosed by dashed lines) of neuromast (asterisks) during Mtz treatment. (E) A cartoon shows the absence of neuromast with undamaged Schwann cells and lateral nerve. (F) Apoptotic cells were clearly labeled by the TUNEL staining in the Mtz-treated larva and the lateral nerve (arrow) stayed undamaged. (G) Mtz was used to treat larvae from the cross of *Tg(foxD3:GFP)* and *Tg(-8.0cldnb:NTR-hkiKGR)*. It appeared that Schwann cells (arrowhead pointed green rod) appeared intact in compared to the disintegrating neuromast (asterisk) in a representative stacked image. The faint broad green signal underneath is stacked signals from underlying tissues. (H-J) After washing out Mtz, most of interneuromast cells (30/34) in the proximity of ablated neuromasts did not form a new neuromast. (K-M) A few (2/34) larvae had regenerated a neuromast (M) through cluster formation (4/34) (L). Scale bars are the same in all panels but only shown in (B).

We thus incubated 3-dpf *Tg(HG7L; -8.0cldnb:NTR-hKikGR)* larvae with Mtz for 12 hours and then washed out to see whether the ablated L3 neuromast could be regenerated. We observed the gap of the missing neuromast was filled, but most fish (30/34) did not regenerate the L3 neuromast even at 4 days after the washout of Mtz (Fig. 5H-J). However, a few larvae (4/34) still formed a cluster at 2 days post Mtz washout (Fig. 5L) and two of them (2/34) even regenerated neuromasts at 4 days post washout (Fig. 5M). This suggests other factors might override the inhibition from pLLn/SWC to activate INCs.

### Macrophages relieve the neural inhibition on interneuromast cells in a cytokine-independent manner

Leukocyte infiltration is the first line of innate immune response. We hypothesized that infiltration of leukocytes may be involved in the regulation of neuromast regeneration. Using *Tg(mpx:eGFP)* larvae, we observed that neutrophil recruitment reached a peak at 6 hours and declining within one day post injury. The neutrophil recruitment could be effectively inhibited by diclofenac, a cyclooxygenease (COX) inhibitor [48, 70] (Fig. S4). We first inhibited the neutrophils recruitment by treating tail-amputated 3-dpf larvae with diclofenac, but still observed a similar regime of neuromast regeneration (Fig. S5A). We further eliminated macrophages with the injection of control or clodronate-loaded liposome (clodrosome), which induce apoptosis of phagocytes [48, 71], into blood circulation via the heart of a 3-dpf larva before tail amputation. Interestingly, regeneration process was slowed down with less than half of larvae (11.5 %) regenerating a new neuromast at 4 dpa in the absence of macrophages (Fig. S5B), supporting the involvement of macrophages in neuromast regeneration upon tail amputation.

To test whether macrophages are involved in the residual regeneration capacity seen in Mtz-treated larvae retaining strong inhibition from intact SWCs, we treated larvae with clodrosome and found significantly hampered neuromast regeneration by inhibiting macrophages (Fig. 6H). This suggests an independent role of macrophages in alleviating the inhibition signals from pLLn. Using larvae from the cross of *Tg(HG7L; -8.0cldnb:NTR-hKikGR)* and *Tg(mpeg1:mCherry)*, a macrophage reporter line [51], we observed that some resident macrophages originally positioned around neuromast (Fig. 6A), then were recruited to the injured neuromast in a polarized and lobulated morphology (Fig. 6B-D). Polarization status has been reported as important features of M1 or M2 macrophages [36]. The M1 macrophages act as pro-inflammatory cells to engulf apoptotic cells (indicated by arrowheads, as green signals within red cells). In a later phase, M2 macrophages, transformed from the M1 status, display dendritic morphology and are essential for inflammation resolution (anti-inflammatory) and tissue modeling. At 2 dpMtz, many elongated macrophages were still seen to interact with the injured pLL system compared to only a few neutrophils (yellow asterisk) existed in larvae from the cross of *Tg(mpeg1:mCherry; mpx:GFP)* [52] and *Tg(HG7L; -8.0cldnb:NTR-hKikGR)* (Fig. 6E-G).

**Figure 6.**
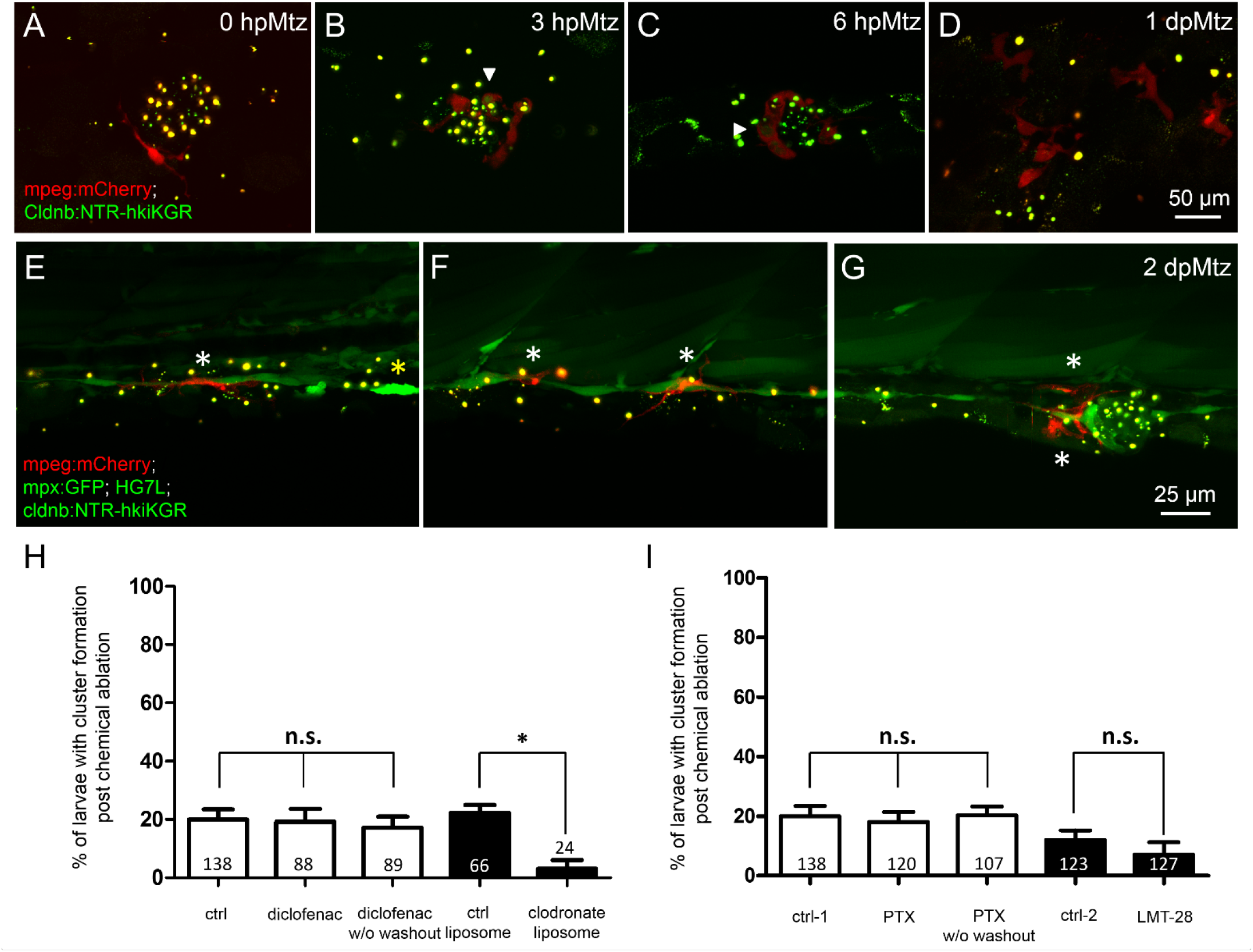
Changes in phagocytes surrounding deteriorating neuromasts and their effects on cluster formation. Larvae from the cross of *Tg(mpeg:mcherry)* (red) and *Tg(-8.0cldnb:NTR-hkiKGR;myl7:EGFP)* (green) were treated with metronidazole (Mtz) as described in Fig. 5, examined and photographed at designated hours or days post Mtz treatment (hpMtz or dpMtz). We observed the disintegration of neuromasts post Mtz treatment (A-D). Red macrophages (arrowheads) were recruited to engulf injured neuromasts in a dendritic cell-like shape within hours post Mtz treatment (A-C). The injured neuromast was disappeared, but macrophages were still retained in the injured site at 1dpMtz. Please note that the macrophages became more compact round shape with protrusions (D). (E-G) Triple-transgenic larvae as designated were treated as described above, examined and photographed at 2 dpMz. Macrophages (red, white asterisks) were still found in the posterior lateral line system, even in the presence of a new neuromast as shown in (G). In contrast, only a few neutrophils (bright green, yellow asterisks) were found. Scales are the same for each row, but only shown in the far-right panel. (H) Larvae from the cross of *Et(HG7L) and Tg(-8.0cldnb:NTR-hKikGR;myl7:EGFP)* were treated without (ctrl) and with 3 μM diclofenac (with or without washout), control or clodronate liposome (H), or treated with different cytokine inhibitors PTX (35 μM) or LMT-28 (10 μM) (I). The above treated larvae were undergone a 12-h Mtz treatment and scored for the % of larvae with cluster formation. Data represents mean ± s.e.m. and analyzed by one-way ANOVA, \**P* < 0.05. n.s.: not significant.

Recently, accumulating evidences point to pivotal role of macrophages in wound repair and tissue regeneration mainly via secreted cytokines [72]. We thus blocked several candidate cytokines secreted by macrophages such as *tumor necrosis factor alpha* (TNFα) and *interleukin-6* (IL-6) with pharmacological drugs including pentoxifylline (PTX) shown to inhibit *tnfα* transcription [73–75] and LMT-28 targeting the IL-6 receptor β subunit (glycoprotein 130, gp130) [76]. Compared to the designated DMSO or methanol control, neither PTX nor LMT-28 treatment ruined the enduring capacity for neuromast regeneration (Fig. 6I). These results suggest that macrophages may exert their effect on neuromast regeneration in a cytokine-independent manner.

### Macrophages patrol in between injury site more often in later regeneration phase

To further understand the interaction between macrophages and regenerating neuromasts, we aimed to dissect more thoroughly their spatial and temporal relationship. First, the larvae from the cross of *Tg(mpeg1:mCherry; FoxD3:GFP)* and *Tg(-8.0cldnb:NTR-hKikGR); Et(HG7L))* were treated with Mtz and examined under LSFM. Larvae were monitored in a wide-field of view (1.3 mm X 1.3 mm), focusing on neuromast L3 to L5, with about 200 z-sections (step size of 1 μm) every five minutes (Fig. 7A). For analytic convenience, z sections were stacked by maximum intensity projection and then region of interest (ROI) was cropped for further image processing. We used the plugin “TrackMate” in Fiji software [61] to analyze mCherry-marked macrophages under red channel with manual corrections of spot labeling to indicate the center of a macrophage and spot linkage to show the migrating route of a macrophage at two continuing time points (Fig. 7A). We also excluded macrophages which did not in contact with the fluorescent pLL from the HG7L line by built-in filters. Every tracking path of individual macrophage could thus be presented by its index or displacement with color map (Fig. 7A, right). By recording the location of each macrophage in different time points (x_n_, y_n_, t_n_) we could acquire useful features to distinguish specific behaviors (Fig. 7B). For example, displacement against distance could be calculated by either the difference between the final and initial positions or the sum of every movement. Then, the velocity or speed of individual macrophage could be calculated.

**Figure 7.**
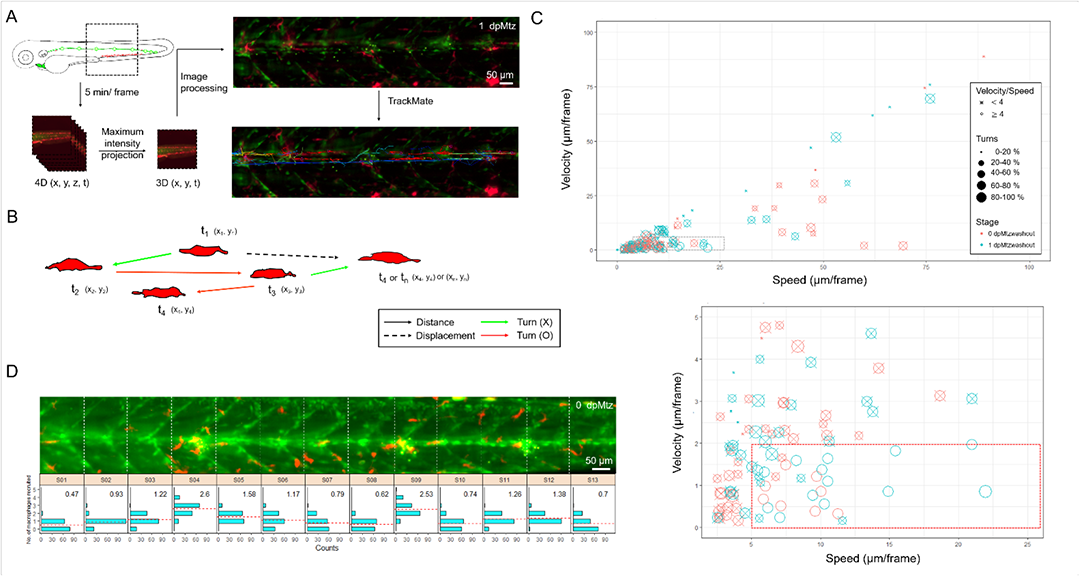
In toto imaging analysis reveals differential cell behaviors and specific spatial distribution of macrophages during neuromast regeneration. (A) As shown in the flowchart, the trunk region including neuromast L3-5 were imaged *in toto* by light-sheet fluorescence microscopy (LSFM) to monitor macrophage behavior during neuromast regeneration. Larvae from the cross of *Tg(mpeg1:mCherry; FoxD3:GFP)* and *Tg(-8.0cldnb:NTR-hKikGR); Et(HG7L)*, a wide field of z-stack images focusing around the cloaca (dashed rectangle) was taken every 5 min. We then reduced the order to 3D (x,y,t) with stacking of z sections and processed to have a better resolution by subtracting background. Finally, we used the plugin “TrackMate” in the Fiji software and manually modified to acquire the tracking path of every macrophages (shown in right-bottom corner). (B) Distance (arrows) and displacement (arrows with dashed line) were measured to calculate speed and velocity as shown in (C). Turns were counted among cell movements (indicated by red) to generate the ratio of turns in (C). (C) A scatter plot shows specific speed (the X-axis) and velocity (the Y-axis) for each macrophage. Other features such as “Speed / Velocity”, “Ratio of turns” and “Stage” are also depicted as shown in the legends on the right to better distinguish different cell behaviors. A region with a speed lower than 25 and velocity smaller than 10 (dashed rectangle) was magnified as shown in the lower panel. (D) Histograms divided by sections (width of 100 μm) represent counts (the X-axis) of different number of recruited macrophages (the Y-axis) during neuromast regeneration. In each panel, the mean of “number of recruited macrophages” is shown in right-upper panel and presented with dashed lines in red.

As shown in the Supplementary Video 5, different cell dynamics of macrophages exist during regeneration. Therefore, we analyzed movies from two larvae each at 0 and 10 hours post Mtz washout. As expected, some macrophages resided at the injury site for a long while (Fig. S6, arrows). In contrast, some macrophages patrolled across the injured site at different range (Fig. S6, arrowheads) and a few of them passed by rapidly (Fig. S6, asterisks). The “patrol” behavior is characteristic of high speed-velocity ratio and frequent turns of direction (Fig. 7B, red arrows). To have a better view of different cell behaviors exhibited by all macrophages in the whole-trunk region of Mtz-treated larvae, the scatter plot was presented in Fig. 7C. Each circle represents one macrophage. More than half of macrophages moved gently and slow (0.2∼2 μm/min). In contrast, a population of actively moving macrophages (Speed: > 1 μm/min, Velocity: < 0.4 μm/min, Speed/Velocity > 4, Ratio of turns > 40%) increased significantly in later regenerative phase at 1 day post Mtz washout (Fig. 7C, marked by a red rectangle in dashed line at the bottom). For a better visualization, other macrophages out of this group are marked out with a ‘X’. This increase in active macrophage migration might be relevant to neuromast regeneration.

To further quantify the distribution of macrophages along the pLL system, we videotaped a Mtz-treated larva at 5 minutes per frame from 0-12 h post Mtz washout (Supplementary Video 6). For analysis, we divided the trunk into 13 sections (S01-S13, each with 100 μm in width) according to the chevron-shaped muscle segment which is clearly marked in the *Et(HG7L)* line. We counted the number of recruited macrophages in each section at each frame and presented a histogram to show the distribution and average numbers of macrophage accumulated from all frames (Fig. 7D). It appeared that more macrophages were recruited within sections containing injury sites (S04, S09, S11 and S12). Unexpectedly, uneven distribution of macrophages was found in sections without injury sites (S5: 1.58 & S6: 1.11 vs. S7: 0.79 & S8: 0.62). This clearly indicates the injury-induced recruitment of macrophages and implies a possible involvement of macrophage in neuromast regeneration. Altogether, we hypothesize that the more frequent macrophage patrolling around injured neuromasts in later phase of regeneration may account for the residual regeneration capacity in Mtz-treated larvae.

### Macrophage intervention may lift the neural inhibition of interneuromast cells

By close examination of those *in toto* time-lapse movies through z-axis, we found macrophages were situated from the top surface to the bottom of trunk (Supplementary Video 7). Macrophages often crawled on INCs with underneath SWCs in close proximity. Interestingly, we observed that some macrophages protrude into the limited space in between INCs and SWCs. To go more in detail, we thus examined this hierarchical structure under confocal microscopy at a higher magnification using a 63X oil lens to dissect the positions of macrophage, INCs and SWCs along the Z-axis and further clarification in orthogonal views (Fig. 8A-C). Normally, macrophages lied on top of INCs and SWCs as shown in Fig. 8A-B. However, macrophages could squeeze into the tiny space between INCs and SWCs as shown in Fig. 8C. It implies that the macrophage may interfere with the physical contact between INC and SWC.

**Figure 8.**
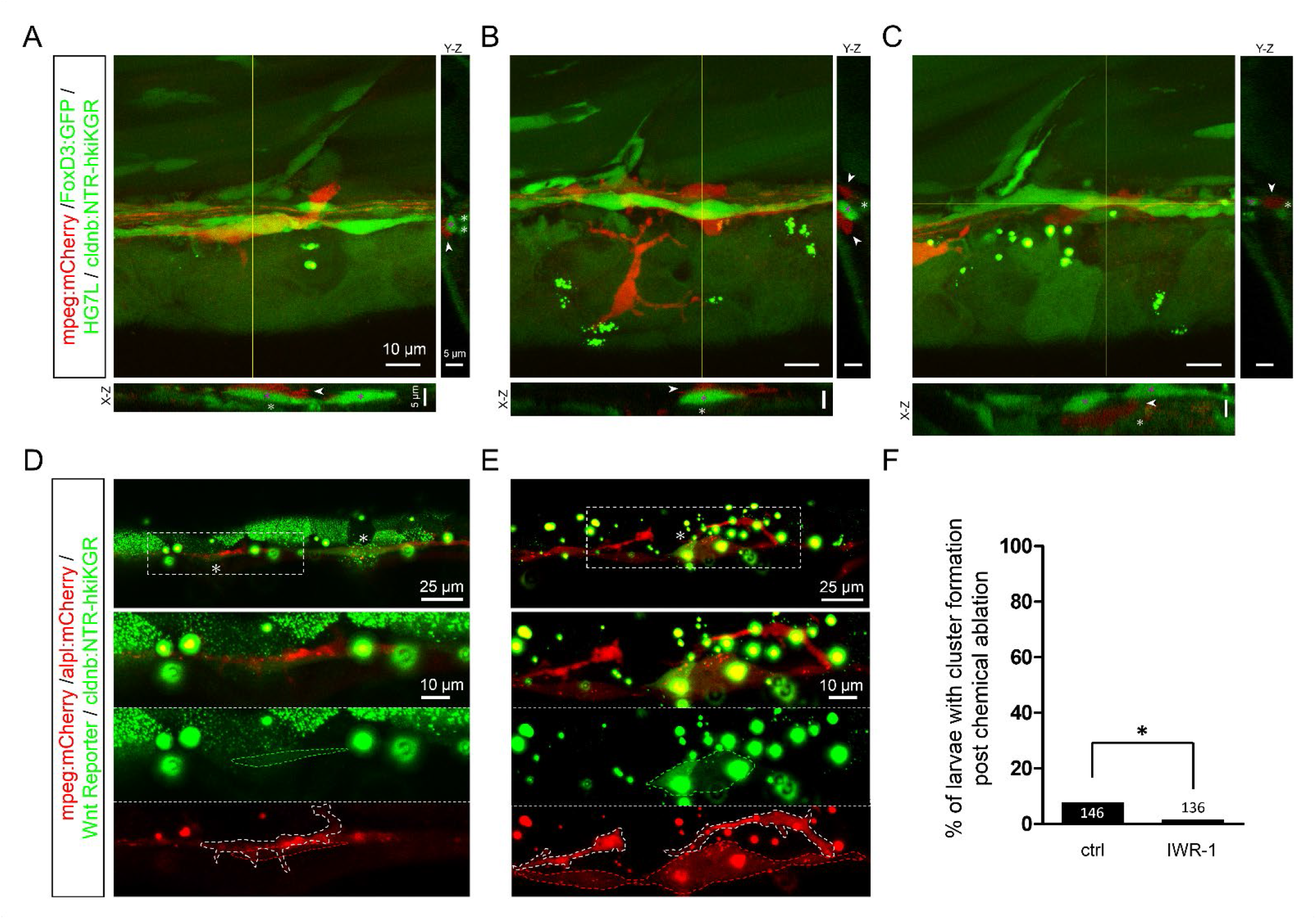
Infiltrated macrophages are found in between Schwann cells and interneuromast cells with higher Wnt activity in Mtz-treated larvae. (A-C) Images shown are representative confocal stacked images of Mtz-treated larvae from the cross of *Tg(mpeg1:mCherry; FoxD3:GFP)* and *Tg(-8.0cldnb:NTR-hKikGR; Et(HG7L))* in orthogonal views, while the X-Z and Y-Z views at the position of yellow lines are shown below and on the right side, respectively. Macrophages (red) indicated by arrowheads were mostly crawling on interneuromast cells (INCs, brighter green, marked by magenta asterisks) in (A-B). In some cases, macrophages were found in between INCs and Schwann cells (SWCs, dim green, labeled by white asterisks). Scale bars are the same but only provided in (A) for respective images. (D-E) Images presented are representative stacked confocal images of Mtz-treated larvae from the cross of *Tg(mpeg1:mCherry; -8.0cldnb:NTR-hKikGR)* and *Tg(-4.7alpl:mCherry; 6XTcf/LefBS-miniP:d2GFP).* Areas shown higher Wnt activity are boxed with dash lines and indicated by asterisks. Magnified diagrams for the dashed rectangles are shown in different color channels to reveal INCs (red, top panel), Wnt signal (dim green, dashed region, middle panel) and macrophage (red, dashed region, bottom panel). Interestingly, some macrophages (enclosed with white dashed line) were found to interact with these activated INCs. Scale bar goes to 25 μm or 10 μm in magnified diagrams. (F) Larvae as above were treated without (ctrl) or with IWR-1 to disrupt Wnt activity and the percentages of larvae with cluster formation were counted as described previously. Data represents mean ± s.e.m. and analyzed by one-way ANOVA, \**P* < 0.05.

Upon the second lateral line development, the INCs on the first lateral line are pushed ventrally by the migrating 2^nd^ primordium to be far away from the inhibitory signal of SWCs. This activates the quiescent INCs to form intercalary neuromasts. It suggests the close relationship between INCs and SWCs is critical to keep INCs silent. We thus hypothesized that the macrophage infiltration could step in to block the neural inhibition. To test this, we first analyzed whether Wnt activity is upregulated in the Mtz-treated larvae. Indeed, increased Wnt activity was seen in lateral line cells or cell clusters during regeneration in Mtz-treated larvae from the cross of *Tg(mpeg1:mCherry; -8.0cldnb:NTR-hKikGR)* and *Tg(-4.7alpl:mCherry; 6XTcf/LefBS-miniP:d2GFP)* (Fig. 8D-E). Interestingly, some macrophages (enclosed in white dashed line) were observed in close physical contact with INCs (enclosed in red dashed line) with upregulated Wnt activity (enclosed in green dashed line) (Fig. 8D-E, lower panels). Furthermore, the inhibition of Wnt signaling by IWR-1, a β-catenin complex stabilizer [77], the residual regeneration capacity were almost completely abolished (Fig. 8F), implying the effect of macrophages works through the activation of INCs with Wnt signaling.

## DISCUSSION

The keys to unravel the mystery of regeneration are the identification of progenitor cells complementing the loss of organs and the underlying induction mechanism therein. In this study, we removed the L6-8 lateral line neuromast in zebrafish larvae by tail amputation or genetic-chemical ablation of the L3 neuromast, and found unequivocally the nearby INCs can be activated to form a new neuromast. The activation of INCs is at least in part by alleviating the inhibition from pLLn/SWC via the intervening infiltration of macrophages.

We observed three sequential phases constituting the whole regeneration morphogenesis of neuromast. Similar morphogenesis sequence, from filling the gap (Phase I), cell clustering (Phase II) to neuromast regeneration (Phase III) had been reported by electroablation of neuromasts [31, 78]. Our data are in agreement with Sanchez et al. that INCs are multipotent progenitors for neuromasts by indirect labeling or cell transplantation [31]. Furthermore, EGFR signaling has been shown to be unequivocal factor for keeping the quiescent status of INCs since perfect regeneration could be achieved by inhibiting EGFR signaling with AG1478 [31]. However, we observed no gap filling activity before one day post injury that is in contrast to the work by Sánchez et al. [31]. It is possible that electroablation might cause an instant injury of the underlying SWCs and pLLn. As in our case the diminishing signal of pLLn in the nearby region of cutting site might indicate a lower expression of Neuregulin 1 type III (Nrg1-3) which is involved in the migration, proliferation and differentiation of SWCs [27] via the receptor, ErbB2 or ErbB3, of SWCs [25, 79]. This tripartite relationship is well established to explain precocious intercalary neuromast formation with the interruption of this ErbB/Neuregulin pathway [26]. Whichever mechanisms they adopted, INCs could respond to neuromast ablation as multipotent progenitors by proliferating and replenishing the loss of organ.

Tissue damage, either sterile or infectious, usually accompanies with innate immunity to protect the organism at the first front line. This inflammatory response features sequential recruitment of different leukocytes to the injury site [34]; neutrophils serves as pioneers while macrophages arrives later on as typical immune process that were seen previously in zebrafish [70, 80, 81] and also evident in our finding (Fig. S4). Neutrophils are capable of eliminating pathogens by phagocytosis and maintaining inflammation with IL1-β secretion [82]. This inflammation is augmented by myeloid cells and surrounding injured cells. They further recruit macrophages to participate in a positive feedback loop. Moreover, macrophages with morphology change from round to dendritic shape were also reconfirmed in our data (Fig. 6A-D) that the polarity transition from M1-like (pro-inflammatory) to M2-like (non-inflammatory) [37] was almost completed within one day post injury. This transition, also known as inflammation resolution, was triggered by the inhibition of IL1-β signaling through TNFα secretion [83] and efferocytosis, a process that activated or infected neutrophils are engulfed by M1-like macrophages [38]. Thus, prolonged inflammation was detrimental since it not only induce more apoptosis but threaten regeneration [84]. Interestingly, M1-like macrophages could promote zebrafish fin regeneration through TNFα/TNFR1 on blastema cell proliferation [41] and stimulate myogenic precursor cells in mammals to divide through secretion of TNFα, IL1-β and IL-6 [85–87]. This promotion likely relies on alleviation of IL1-β signaling, achieving inflammation resolution, instead of cell debris clearance or cell death prevention [83]. On the other hand, as being anti-inflammatory, M2-like polarized macrophages could also be involved in inflammation resolution via secretion of anti-inflammatory molecules such as TGF-β1 or IL-10 and further tissue remodeling [36, 41]. In our hands, the residual regeneration capacity was not affected by inhibiting TNFα signaling, indicating M1-like macrophages may not play an essential role through secretion of TNFα. Nevertheless, we need further investigations like the ablation of macrophages at different time points after injury or the use of transgenic line, *Tg(tnfa:eGFP-F)* to distinguish two classes of macrophages [36], in order to resolve the contribution of M1-like or M2-like macrophages in the neuromast regeneration.

We performed macrophage tracking post injury with wide-field acquisition and categorized individual cell behaviors as “Stay”, “Patrol” and “Flash”. Tracking analyses of immune cells after tissue loss were previously conducted in fin or retina regeneration [36, 70, 81, 88, 89] but carried out with simple parameters such as velocity and speed. With these parameters, cell heterogeneity was established between peripheral and resident macrophages [89], or macrophages behaviors upon different injury condition [88]. We introduced more parameters revealing direction (Velocity/Speed), orientation (Turns) and phase to identify novel cell subclasses. While patrolling macrophages increase in later phase of regeneration (1 dpMtz), all three behaviors appeared in both early (0 dpMtz) and late phase. That means, this novel categories may not specifically correspond to M2-like macrophage which could be further subdivided into M2a, M2b and M2c subclasses in mammals [90]. Furthermore, macrophages seem to stay in hot zones and shuttle in between SWCs and INCs with high z-resolution analysis. Thus, we hypothesize that more certain M2-like macrophages (1 dpMtz) could interrupt the connection between SWCs and INCs in specific region via unknown mechanism. The dissociation ability could be either physical or chemical. Physical force could be generated by macrophages to adhere and connect the ruptured blood vessels [47]. INCs were also witnessed to be pushed away while macrophages passed. Further *in vivo* dynamics of actin filaments regulated by phosphatidylinositol-3-kinase (PI3K) or Ras-related C3 botulinum toxin substrate 1 (Rac1) could possibly help reveal the filopodia or lamellipodia-dependent cellular action. Chemical reaction could be extracellular matrix (ECM) remodeling by matrix metalloproteinases (MMPs) from macrophages [91]. ECM remodeling further mediating leukocyte recruitment was proved to be regulated by critical expression of MMPs family enzyme such as MMP-9 or MMP-13 and be crucial to heart regeneration in zebrafish [92]. Interestingly, since macrophages are supposedly gifted to separate INCs and SWCs, they were found to frequently visit along the migratory trajectory of 2^nd^ pLLp. Whether macrophages promote intercalary neuromast formation via dissociation of SWCs and 1^st^ lateral line needs to be further investigated. Therefore, we hypothesize here that macrophages could play essential role of separating INCs and SWCs both in development and regeneration. In development, while macrophages could digest the linkage between SWCs and 1^st^ lateral line, 2^nd^ primordium could help create enough space physically to alleviate the inhibition from SWCs, resulting robust intercalary neuromast formation. During regeneration, M2-like macrophages could only separate SWCs and INCs by itself. Limited freedom of neural inhibition with minimal gap between INCs and SWCs generated by individual macrophages thus lead to low successful rate of regeneration. Macrophages are recently well-known to participate widely in development, homeostasis and regeneration [35, 93], this the first study to suggest that macrophages could support different scenario, i.e. regeneration and development in zebrafish lateral line, by utilizing similar mechanism.

In conclusion, the inhibition from SWCs and lateral line nerves is the key factor keeping the quiescence of INCs. The quiescence of INCs can be alleviated by fin amputation or possibly via intervening macrophage in between lateral line and SWCs (Fig. 9). In both cases, the regeneration of neuromast can be blocked or delayed by inhibiting macrophages. Macrophages may also be involved in intercalary neuromast formation during development. Altogether, our results strongly suggest that macrophages may participate in the development of neuromast. More importantly, they play a pivotal role in awaking interneuromast cells to regenerate neuromasts in an injured lateral line in zebrafish.

**Figure 9.**
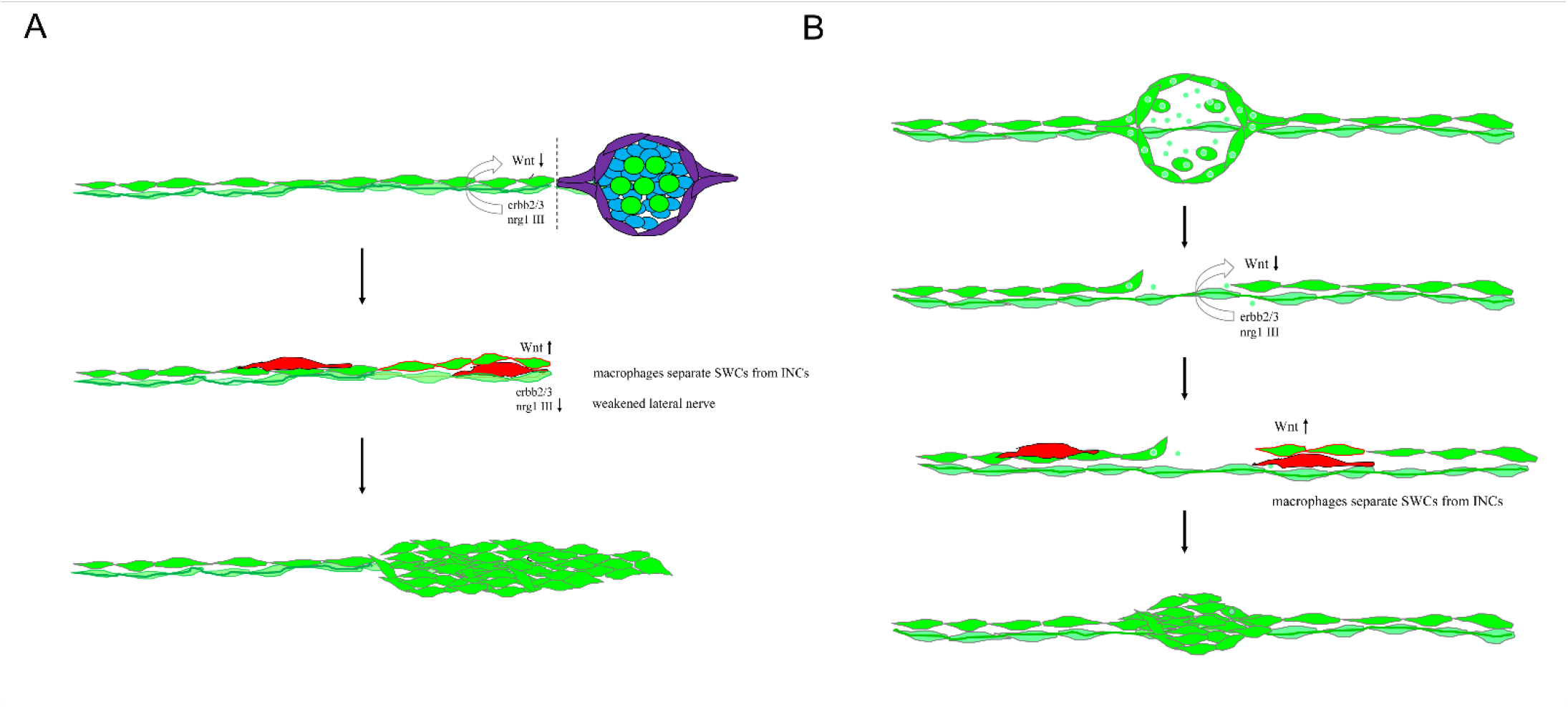
Mechanistic models for lateral line regeneration. (A) Under normal conditions, the pLLn (thick dark green thread) nrg1-III-activated erbb2/3 receptors within Schwann cells (SWCs, light green) keep interneuromast cells, (INCs, bright green) quiescent by limiting the Wnt activity. Upon the loss of distal neuromasts by tail amputation, the lateral line nerve is weakened (thin light green thread) in the proximity of cutting edge to break the quiescence of INCs (red outlines). The Wnt activity in INCs is elevated and results in the formation of cluster. (B) Upon specific ablation of neuromasts by NTR-hKikGR protein (bright green spots)-induced by Mtz without damaging surrounding posterior lateral line nerve and SWCs, INCs remain quiescent by integral EGF inhibition. Most macrophages crawl onto INCs which result in filling the gap without successful neuromast regeneration. However, occasionally macrophages could infiltrate in between INCs and SWCs. This breaks the contact between INCs and SWCs and the inhibition of EGF signaling. INCs are then activated to be able to differentiate and regenerate new neuromasts through cluster formation.

## ACKNOWLEDGEMENTS

We thank Dr. Tatjana Piotrowski from Stowers Institute for Medical Research for providing p5E-*8.0cldnb* plasmid. We thank Dr. Aaron Steinier from Pace University for providing *Tg(-4.7alpl:mCherry)* transgenic fish. We thank Dr. Chung-Der Hsiao from Chung Yuan Christian University in Taiwan for providing pME-NTR-hKikGR plasmid. We thank Taiwan Zebrafish core facility for providing several transgenic lines used in this study. We thank Ms. Shu-Chen Shen in Advanced Optical Microscope Core Facility in Agricultural Biotechnology Research Center in Academia Sinica for great assistance with light sheet microscope. We thank Ms. Yi-Chun Chuang in Technology Commons in National Taiwan University for excellent assistance with confocal microscopy. The authors would also like to express great appreciation to the staffs of the zebrafish Core at National Taiwan University for the assistance in fish maintenance. The work was supported by grants from the Ministry of Science and Technology, Taiwan (MOST 108-2311-B-002-015; MOST 109-2311-B-002-020) to SJL.

## SUPPLEMENTARY FIGURE & LEGENDS

**Figure S1.**
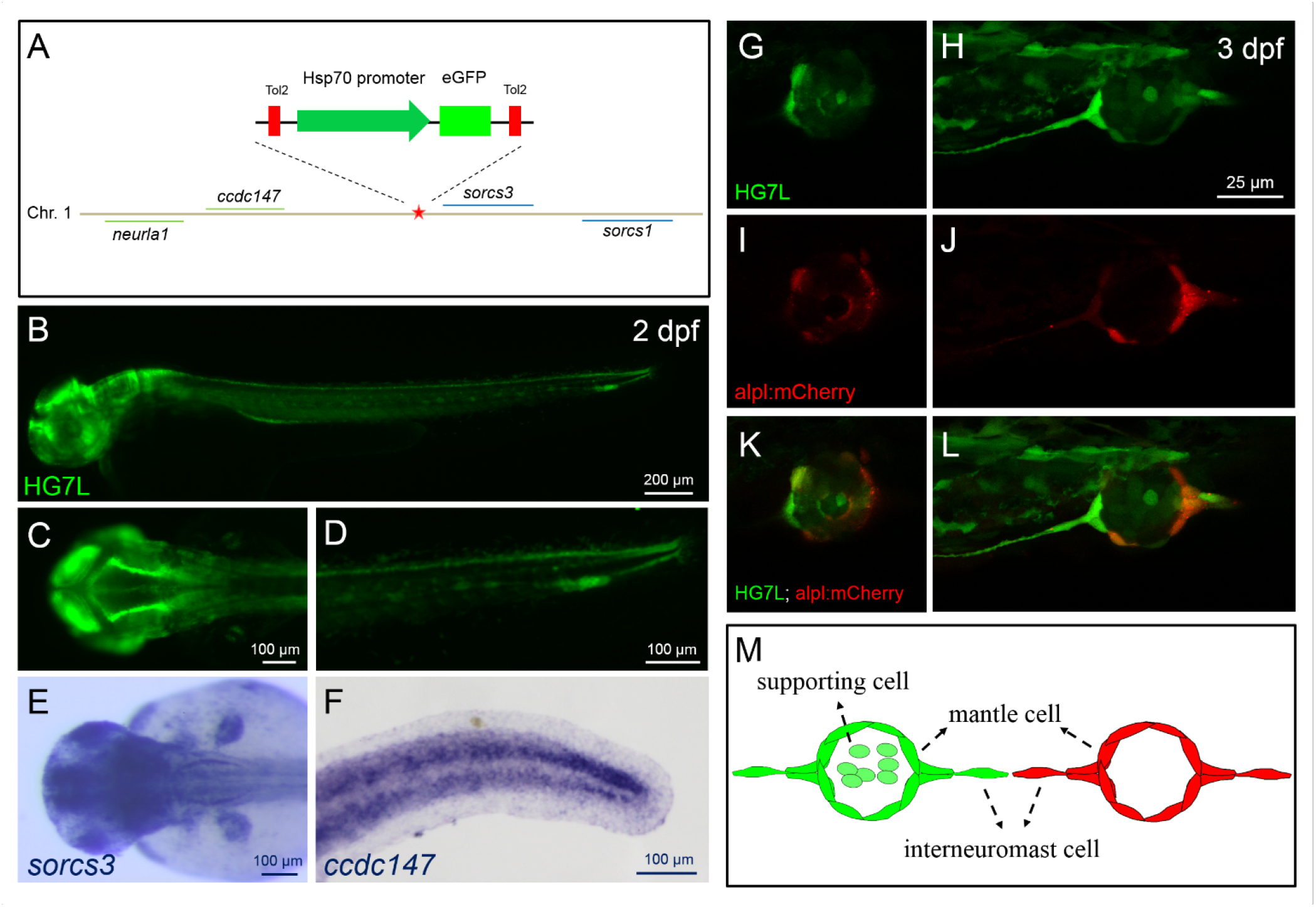
Generation and characterization of an enhancer-trapped line, *Et(HG7L),* expressing EGFP in interneuromast and mantle cells. An enhancer trap line *Et(HG7L*) line was generated by random insertions of a Tol2 construct containing a heat-shock 70 (hsp70) promoter and an EGFP gene. The insertion site was between *ccdc147* and *sorcs3* in Chromosome 1 by inverse PCR analysis as shown in (A). A larva at 2 days post fertilization (dpf) was examined under epifluorescencent microscopy, photographed with anterior to the left in the lateral side (B). Magnified dorsal and lateral view of anterior head and posterior region (D) are shown in (C) and (D), respectively. Larvae were fixed at 2 dpf and subjected whole-mount *in situ* hybridization against *sorcs3* (E, anterior head region in dorsal view) or *ccdc147* (F, posterior tail region in lateral view). (G-L) Larvae at 3-dpf from the cross of *Et(HG7L)* and *Tg(-4.7alpl:mCherry)* were immobilized and examined under confocal microscopy at EGFP (green) and mCherry (red) channel. At the EGFP channel, the *Et(HG7L)* revealed EGFP in interneuromast cells (INCs) and mantle cells (MCs) and also a subpopulation of supporting cells showing weaker expression (G,H). At the mCherry channel, the *Tg(-4.7alpl:mCherry)* had mCherry in INCs and MC, but not supporting cells (I,J). Green and red channel superimposed images are shown below (K,L). (M) A cartoon depicts the expressions of EGFP (green) and mCherry (red) in the *Et(HG7L)* (left) and *Tg(-4.7alpl:mCherry)* (right) lines, respectively.

**Figure S2.**
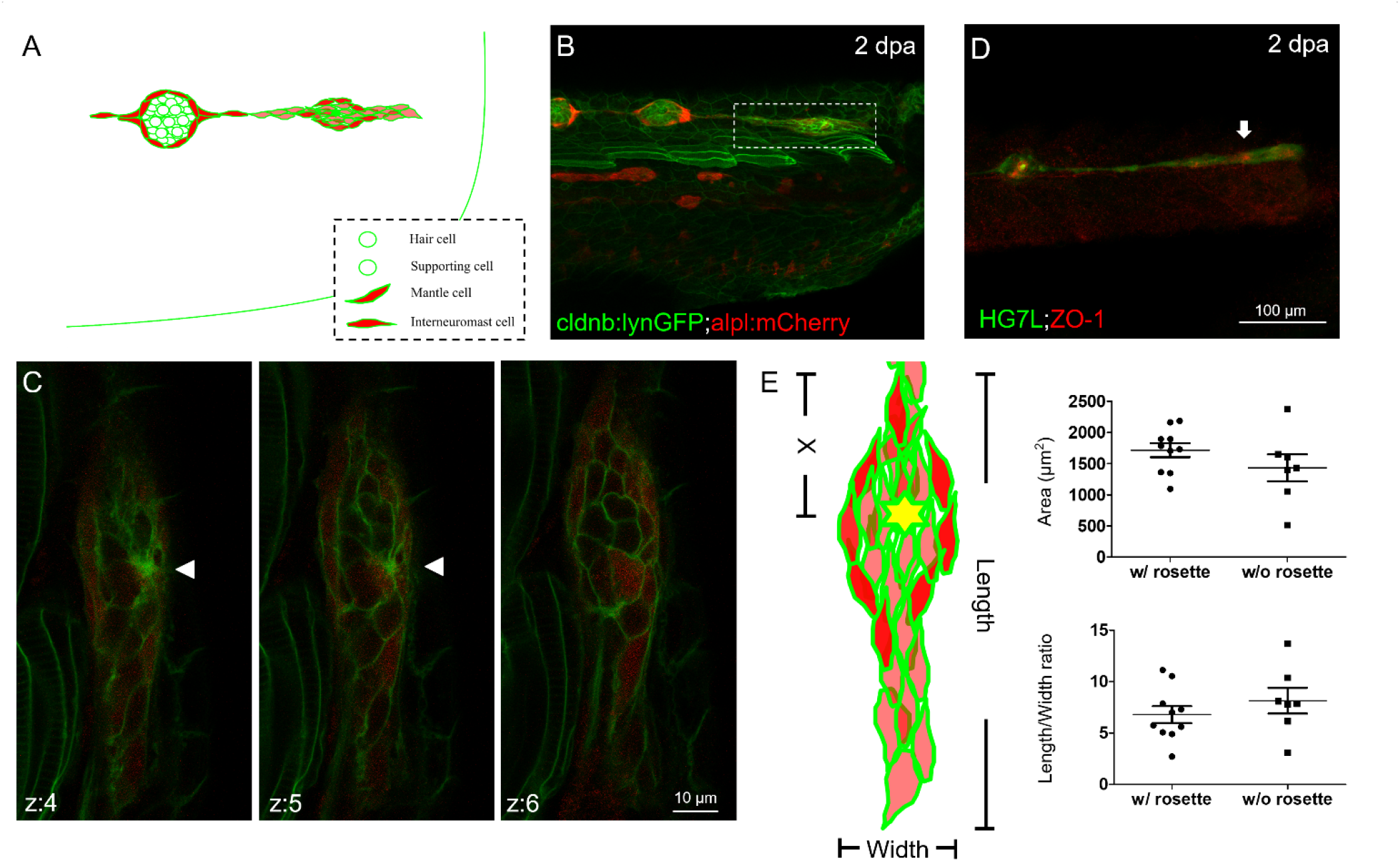
A rosette forms within the regenerating cluster as a landmark of neuromast formation. (A) We used larvae from the cross of *Tg(-8.0cldnb:lyn-GFP) (cldnb:lynGFP in green)* and *Tg(-4.7alpl:mCherry)* (alpl:mCherry in red) to reveal morphogenesis of neuromast regeneration post fin amputation. The cldnb:lynGFP line expressed green EGFP in all cell types of lateral line and the alpl:mCherry line expressed red mCherry in interneuromast cells and mantle cells as shown in the cartoon. (B) At 2 days post amputation (dpa), the double transgenic larvae had formed a cluster (boxed with a dash line) as examined under confocal microscopy. (C) The regenerating cluster was scanned at a higher magnification for 10 1.76-um stacks labeled z1-10. Stack z4-6 are shown and a rosette (arrowhead) was clearly seen in the z-4 and z-5 stacks. (D) Larvae were also fixed and subjected to immunohistochemistry against ZO-1 expressed in tight junctions (red) and GFP expressed in lateral line (green). A strong ZO-1 signal was observed in the cluster as indicated by an arrow that is a sign of polarity establishment. (E) To compare clusters with and without a rosette, we measured the area, length and width of a cluster as shown in a cartoon on the left (a rosette is represented by a yellow asterisk) and calculated their area and length/width ratio. The “X” indicates the distance between the rosette center and the lagging end of cluster. Scatter plots are presented and each dot represents the measurement from one larva. Mean ± s.e.m. are shown. Data were analyzed by one-way ANOVA (n = 18).

**Figure S3.**
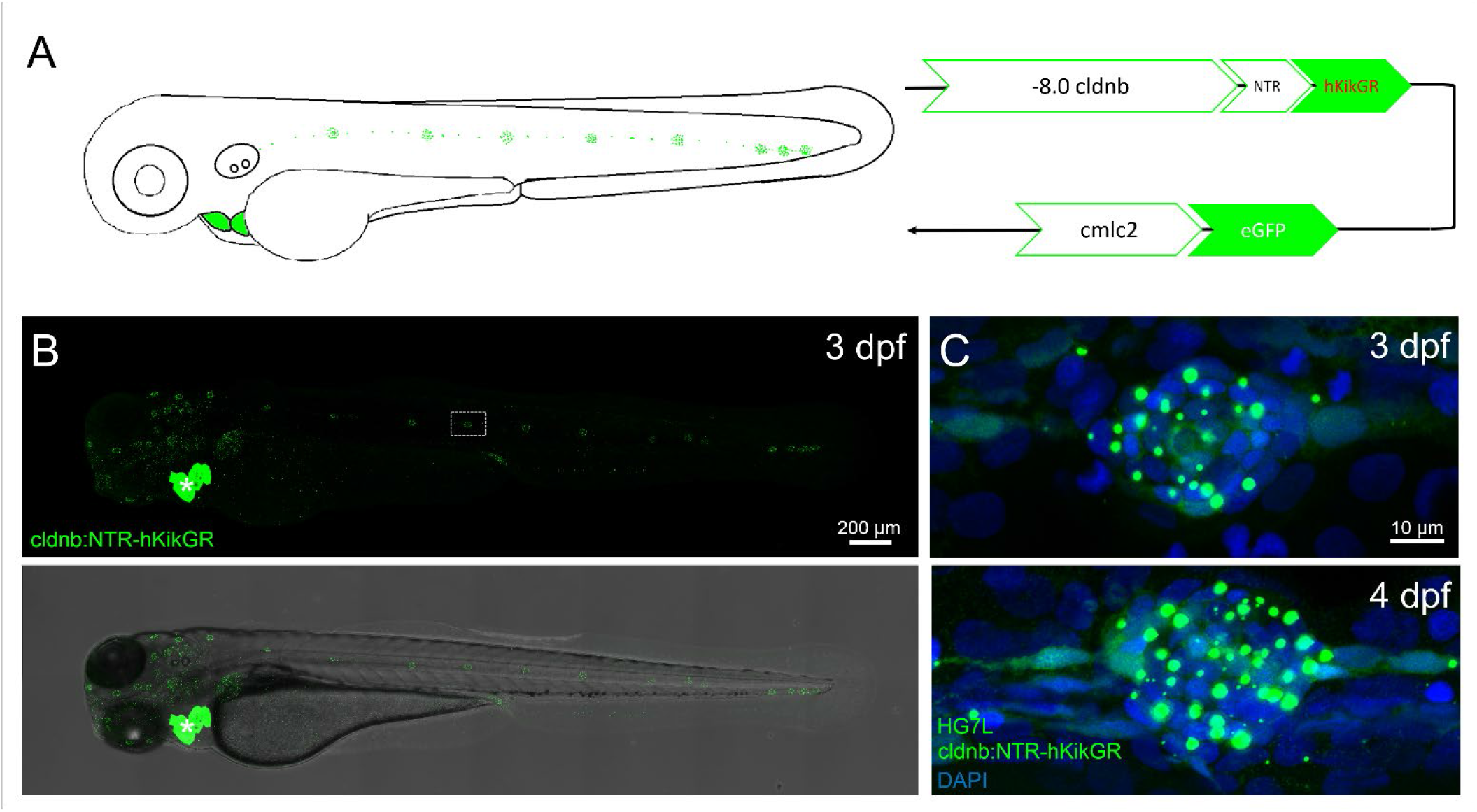
Characterization of a Tg(-8.0cldnb:NTR-hKikGR; myl7:EGFP) double-transgenic zebrafish line. (A) Design of a transgenic cassette composed of an 8-kb *claudin b* promoter (*-8.0 cldnb*), a nitoreductase gene (NTR) fused with a hKikGR gene, a *myl7* promoter and a EGFP gene. The transgenic cassette is expected to express green fluorescence along the lateral line and heart (missing) as depicted in a cartoon on the right. (B) The *Tg(-8.0cldnb:NTR-hKikGR; myl7:EGFP)* larvae at 3 days post fertilization (dpf) were fixed and examined under confocal microscopy at GFP channel. It showed a strong *myl7*-driven green fluorescence in the heart (white asterisk) that served as a convenient selection marker. Punctate green fluorescence was found in neuromasts as shown in a representative neuromast enclosed by a dashed rectangle along anterior (head region to the left) and posterior lateral line systems (trunk region to the right). A superimposed dark and bright field image is shown below. (C) Larvae from the cross of *Tg(-8.0cldnb:NTR-hKikGR; myl7:EGFP)* and *Et(HG7L)* were fixed, incubated with DAPI, examined and photographed at the GFP and UV channel. Representative superimposed images for a neuromast of 3 and 4-dpf larvae are presented. Punctate green fluorescence from NTR-hKikGR proteins was found around blue nuclei stained by DAPI in every cell within a neuromast.

**Figure S4.**
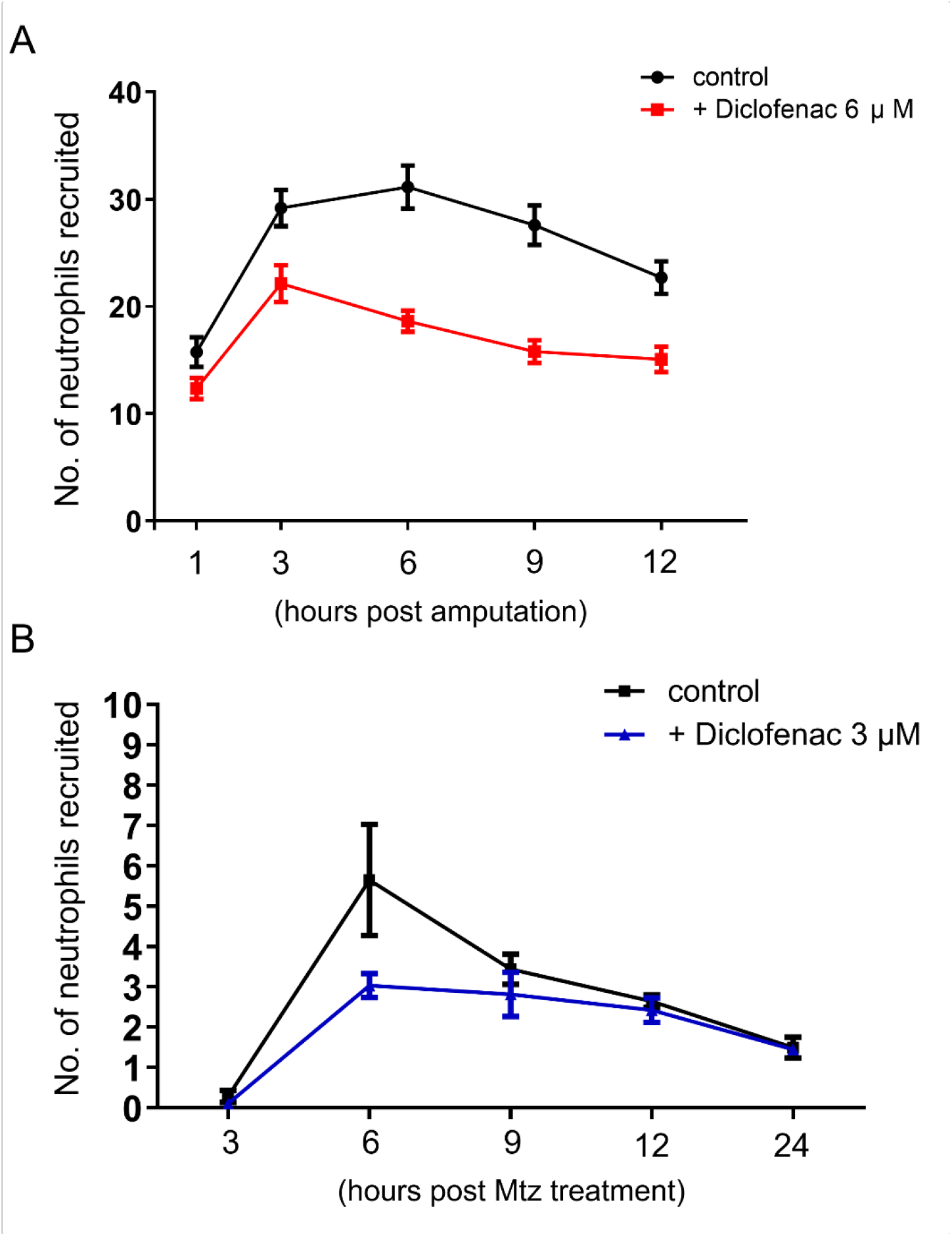
Neutrophils recruitment was impaired by diclofenac sodium post injury. (A) The 3-days post fertilization (dpf) larvae of *Tg(mpx:GFP; Et(HG7L))* with neutrophils labeled by GFP were tail-clipped to ablate neuromasts. After amputation, larvae were transferred to dish without or with Diclofenac and the number of neutrophils recruited to the proximal amputated tail were counted at designated stages (N = 3, n = 25 for the control group, n = 24 for the Diclofenac 3 μM group, n = 27 for the Diclofenac 6 μM group). (B) Mtz-treated *Tg(-8.0cldnb:NTR-hKikGR); Et(HG7L))* larvae were also transferred to dish without or with 3 μM Diclofenac and the number of neutrophils recruited to the injury site were counted at designated stages (N = 3, n = 27 for the control group, n = 31 for the Diclofenac group).

**Figure S5.**
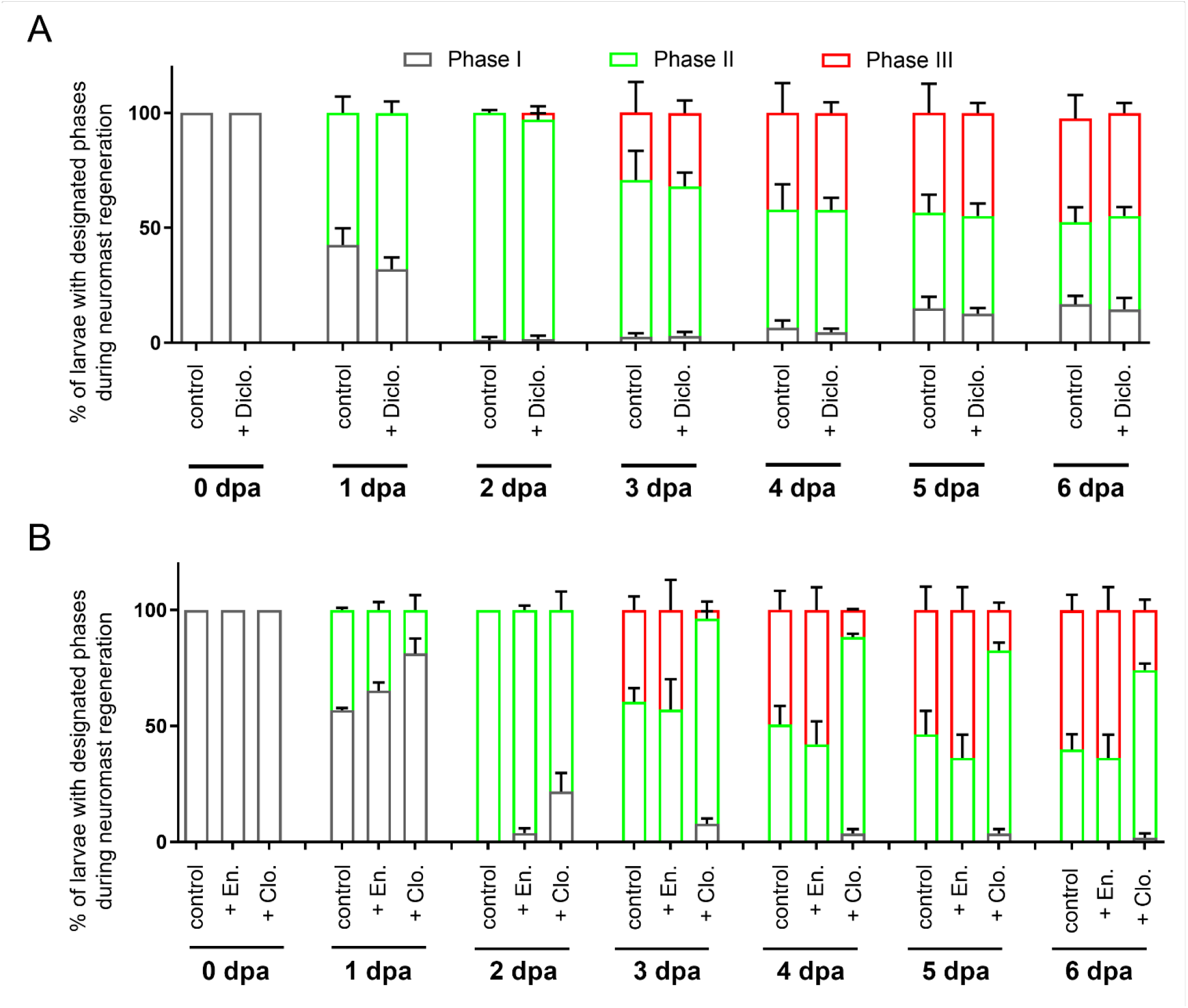
Macrophage ablation retards the regeneration of neuromast post tail amputation. (A) *Et(HG7L)* larvae at 3 day post fertilization were fin amputated and cultured in designated concentration of diclofenac sodium. The number of survived larvae at each phase and treatment were counted at designated stages and the percentages of larvae are shown (N = 4). (B) The vehicle control, encapsome (En), or clodronate liposomes (Clo) were injected into the posterior cardinal vein (PCV) of an *Et(HG7L)* larvae at 3 day post fertilization and the fin was amputated after injection. The numbers of survived larvae at each phase and treatment were counted at designated day post amputation (dpa) and the percentages of larvae are shown (N = 3).

**Figure S6.**
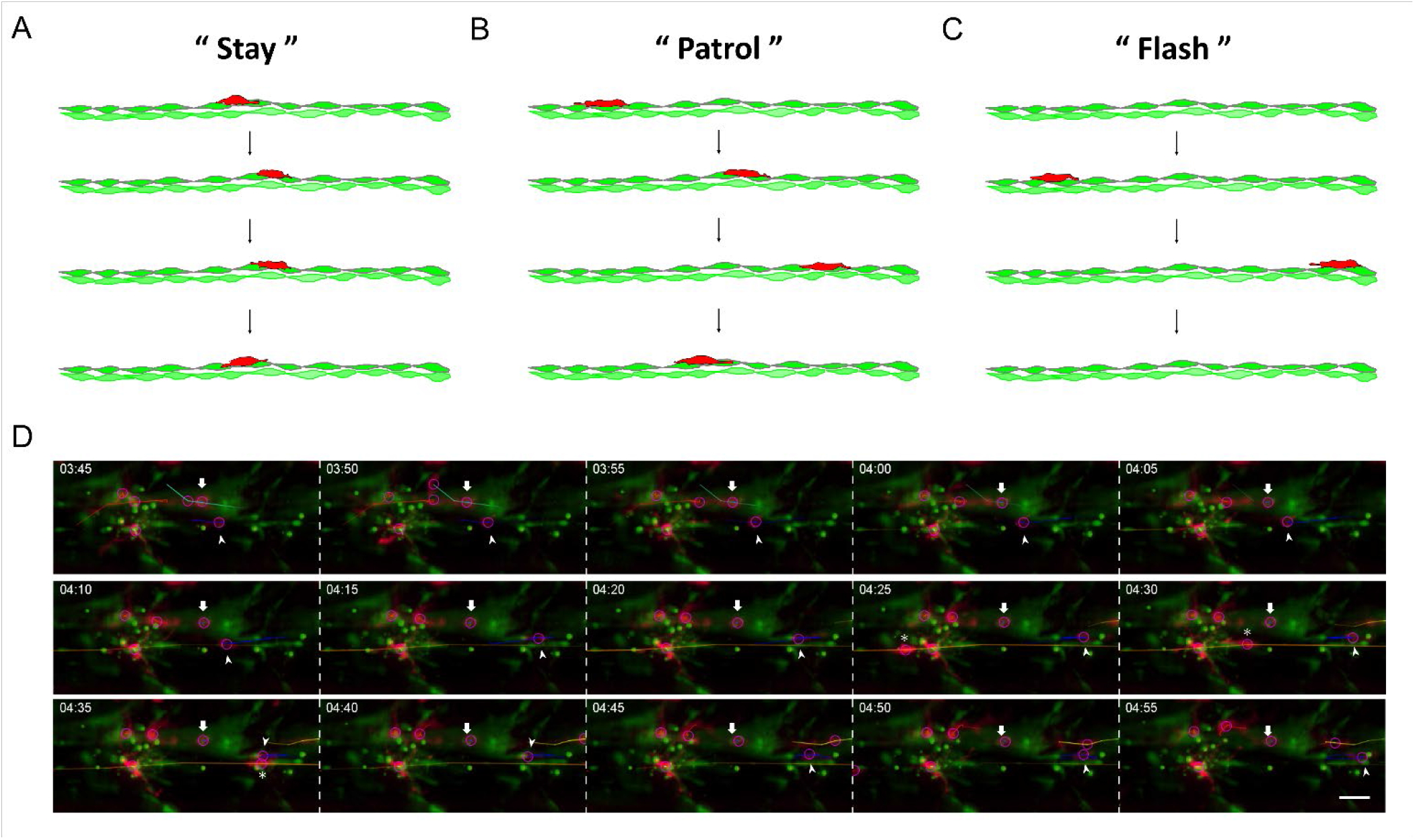
Macrophages displays distinct cell behaviors, including “Stay”, “Patrol” and “Flash” during neuromast regeneration. (A-C) Cartoons show macrophages (red) migrate along INCs (bright green with border line) and underneath SWCs (light green without border line) with differential directions and different velocity. We thus categorized these behaviors as (A) “Stay” with minimal movements, (B) “Patrol” with limited movements and frequent changes in direction, and (C) “Flash” with speedy unidirectional movement. (D) Snapshots of a 70-min movie was cropped from the Video 5. An arrow, arrow head or asterisk is indicating a “Stay”, “Patrol” or “Flash” macrophage, respectively, in each frame. Please note the migrating distance and direction of a macrophage along the X axis to appreciate the difference between different types of behavior.

**Figure S7.**
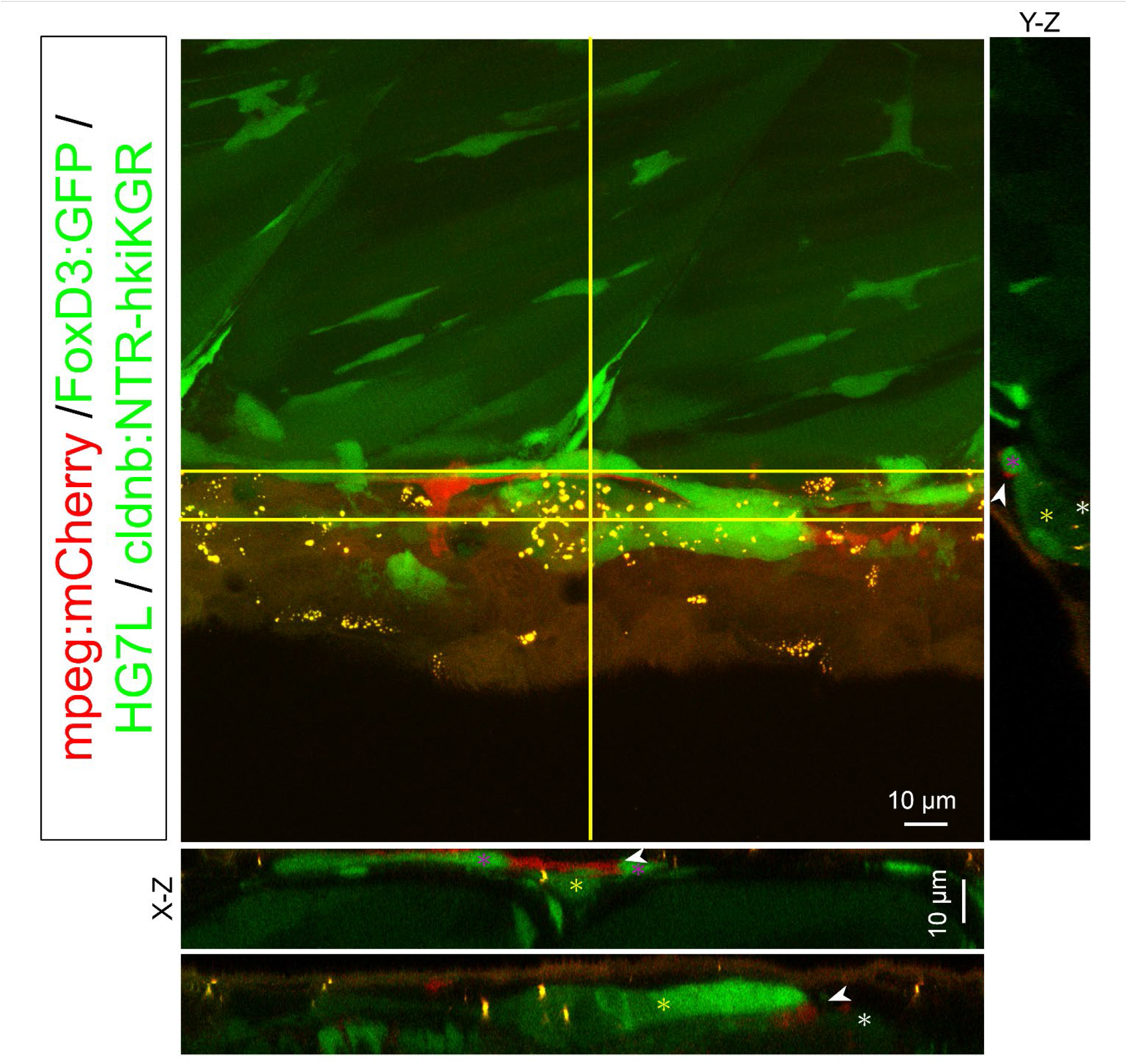
Macrophages interact with 2^nd^ posterior lateral line primordium (2^nd^ pLLp) migration during development. Images shown are representative confocal stacked images of 4-days post fertilization (hpf) larvae from the cross of *Tg(mpeg1:mCherry; FoxD3:GFP)* and *Tg(-8.0cldnb:NTR-hKikGR; Et(HG7L))* in orthogonal views, while two X-Z views and one Y-Z views are shown below and on the right side, respectively. Two separate X-Z views (upper or lower) were indicated by solid or dashed horizontal line. During development, 2^nd^ pLLp (circles in dashed line) would migrate from the otic vesicle at later stages, along the same path of 1^st^ pLLp. The 2^nd^ pLLp would separate SWCs (white asterisks) and INCs, leading to the intercalary formation. Interestingly, macrophages (red) indicated by arrowheads were found in between the 2^nd^ pLLp and INCs or SWCs. Scale bars are the same as 10 μm.

## SUPPLEMENTARY MOVIE CAPTIONS

**Movie S1.** The tail fin of an *Et(HG7L)* larva at 3 days post fertilization (3 dpf) was cut to remove neuromast L6-8 as described in Fig. 1, embedded in agar and imaged under confocal microscopy. Images were taken at 6 min per frame for 6 h. This video shows the Phase I of neuromast regeneration described in Fig. 1. At the leading end toward cutting edge (bottom), most interneuromast cells (INCs) stayed in line with visible cell protrusions (arrowheads). Several INCs (arrows) were seen to crawl onto the original lNC. Time is shown on the top right corner.

**Movie S2.** The *Et(HG7L)* larva was treated described in Video 1. This video shows the Phase II of neuromast regeneration. A large cluster was formed with many visible cell protrusions (arrowheads) at both leading and lagging end toward the cutting edge at the bottom.

**Movie S3.** The *Et(HG7L)* larva was treated described in Video 1. This video shows the Phase III of neuromast regeneration. A new neuromast was formed with a clear ring-like structure and GFP-labeled mantle cells.

**Movie S4.** AG1478 caused hyperactive cell protrusion and cell division. Cell protrusions are marked by arrowheads. Mother cell and two daughter cells are labeled by white and magenta asterisks, respectively.

**Movie S5.** *In toto* imaging analysis shows differential macrophage (red) behaviors during regeneration. Larvae from the cross of *Tg(mpeg1:mCherry; FoxD3:GFP)* and *Tg(-8.0cldnb:NTR-hKikGR); Et(HG7L)* were treated with Mtz and monitored under a light-sheet fluorescent microscope. Time-lapse movies of merged green and red channels (top row), merged channels with cell tracking (purple circles, middle row), and red channel with cell tracking (bottom row) are combined vertically and presented. Images were taken at 5 min per frame for 6 h 35 min as shown on the top left.

**Movie S6.** Uneven distribution of recruited macrophages (red) during regeneration were analyzed and presented as graphic interchange format of bar chart (upper row) to reveal the dynamics of macrophage distribution. Larvae from the cross of *Tg(mpeg1:mCherry; FoxD3:GFP)* and *Tg(-8.0cldnb:NTR-hKikGR); Et(HG7L)* were treated with Mtz and observed under a light-sheet fluorescence microscope. Time-lapse movies of merged channels with cell tracking (middle row), and red channel with cell tracking (bottom row) are combined vertically and shown here. Images were taken at 5 min per frame for 11 h 55 min as shown on the top left of images.

**Movie S7.** This video is a three-dimensional pseudo color reconstruction. While most macrophages (red, left) were crawling on INCs (left), some macrophages (red, middle) could scroll through the space between INCs and underneath SWCs (right).

**Movie S8.** Macrophages (red) could push away interneuromast cells (green, indicated by an arrow) while passing by.

**Movie S9.** Macrophages (red) were not only in contact with interneuromasts (green, arrow) but were also dynamically embracing the second posterior lateral line primordium (green, open arrow) during development.

## Notes

### Competing Interest Statement

The authors have declared no competing interest.

